# A dichotomy in association of core transcription factors and gene regulation during the activated B-cell to plasmablast transition

**DOI:** 10.1101/2019.12.23.884007

**Authors:** Mario Cocco, Matthew A Care, Muna Al-Maskari, Gina Doody, Reuben Tooze

**Author notes:** These authors contributed equally to this work (Mario Cocco, Matthew A Care). Corresponding author: Reuben Tooze, Division of Haematology & Immunology, Wellcome Trust Brenner Building, Leeds Institute of Medical Research @ St James’s, University of Leeds, Leeds, LS9 7TF, UK.

## Abstract

The activated B-cell (ABC) to plasmablast transition is the cusp of antibody secreting cell (ASC) differentiation but is incompletely defined. We apply expression time-courses, parsimonious gene correlation network analysis, and ChIP-seq to explore this in human cells. The transition initiates with input signal loss leading within hours from cell growth dominant programs to enhanced proliferation, accompanied from 24h by ER-stress response, secretory optimization and upregulation of ASC features. Clustering of genomic occupancy for ASC transcription factors (TFs) IRF4, BLIMP1 and XBP1 with CTCF and histone marks defines distinct patterns for each factor in plasmablasts. Integrating TF-associated clusters and modular gene expression identifies a dichotomy: XBP1 and IRF4 significantly link to gene modules induced in plasmablasts, but not to modules of repressed genes, while BLIMP1 links to modules of ABC genes repressed in plasmablasts but is not significantly associated with modules induced in plasmablasts. Pharmacological inhibition of the G9A (EHMT2) histone-methytransferase, a BLIMP1 co-factor that catalyzes repressive H3K9me2 marks, leaves functional ASC differentiation intact but de-represses ABC-state genes. Thus, in human plasmablasts IRF4 and XBP1 emerge as the dominant association with ASC gene expression, while BLIMP1 links to repressed modules with particular focus in repression of the B-cell activation state.

## Introduction

The differentiation of plasma cells (PCs) from B-cells depends on epigenetic reprogramming, which is driven by sequential shifts in transcription factor expression and is division linked.^1, 2^ In the context of B-cell differentiation independent of the germinal centre reaction the intermediate steps consist of the activated B-cell state (ABC), which may also be referred to as a pre-plasmablast,^3^ and the plasmablast, the immediate proliferating precursor of the quiescent PC.^1^ While we understand many elements of these intermediate steps, including key transcriptional regulators and relationships to certain types of mature B-cell neoplasm, details of the activated B-cell to plasmablast transition are limited.

An informative approach to analysis of cell state transitions is the application of time course gene expression data.^4^ This can be used to resolve the sequence of co-regulated gene expression in coordinately responding cells, as observed in PC differentiation.^2^ Analysis with bioinformatics networking tools allows the resolution of fine grained patterns of temporal co-expression across such differentiating cell populations,^5^ which by inference may enrich for common gene regulatory input.^6, 7^ This can be tested against independently derived data such as the genomic occupancy patterns of key transcription factors at specific differentiation states encompassed in the expression time course. While defining the combinatorial logic controlling the expression of individual genes is highly challenging,^7^ associations between transcription factor occupancy and the expression of multiple genes in a co-regulated gene module can allow the identification of shared regulatory enrichment linked to the cell state transitions.^8^

Three transcription factors - IRF4, BLIMP1 and XBP1 - have been principally linked to the reprogramming of gene expression that underpins the ABC to plasmablast transition.^1^ Detailed models have emerged which place these factors along with input activating signals in a hierarchy wherein IRF4 sits downstream of cytokine and NFkB driven signaling pathways, and BLIMP1 downstream of IRF4.^9–11^ BLIMP1 in turn controls XBP1 expression potentially both through transcriptional de-repression, and through ER-stress/unfolded protein response related pathways. The former for example links BLIMP1 to XBP1 upregulation via repression of PAX5,^12^ and the latter links BLIMP1 to the control of immunoglobulin gene transcription and the alternative poly-adenylation switch from membrane bound to secretory forms.^13^ XBP1 is linked to functional secretory optimization, although lack of XBP1 does not preclude phenotypic differentiation.^14–16^ Deletion of each of these TFs in a B-cell specific fashion in mice provides evidence for this broad epistatic relationship. IRF4 deficiency arrests B-cell differentiation prior to the activated B-cell/preplasmablast phase.^9, 10^ BLIMP1 deletion precludes plasmablast/PC differentiation but allows transition to the activated B-cell/preplasmablast phase.^11^ While XBP1 deletion allows the generation of phenotypic PC-like populations which however lack optimization for the secretory function of the equivalent normal population.^14^

A critical step in this process is the expression of transcription factor BLIMP1 (encoded by the gene *PRDM1*).^11^ In the absence of BLIMP1 expression components of the B-cell transcriptional program fail to be repressed, and the reprogramming of PCs for secretory activity is abortive including incomplete metabolic reprogramming and failure to switch from membrane to secretory forms of immunoglobulin.^13, 17^ Such findings in murine genetic models have extended the contribution of BLIMP1 to include a more extensive role in positive regulation of induced gene expression. In human malignancy inactivation of BLIMP1 occurs in a subset of Diffuse Large B-cell Lymphomas sharing many features with the physiological activated B-cell state. In this context loss of BLIMP1 function is predominantly interpreted as failure of BLIMP1 associated repressive functions, which in conjunction with other oncogenic events trap the malignant cells at the ABC to plasmablast transition.^18–20^ However, we know relatively little about the extent to which BLIMP1 couples to either positive or negative regulation of gene expression during the analogous differentiation of primary human B-cells from ABC to plasmablast.

BLIMP1 mediates its role as a transcriptional repressor through the recruitment of epigenetic regulators, which include histone deacetylases HDAC1/2/3, histone methyltransferases G9A (EHMT2) and EZH2, as well as the histone demethylase LSD1, and the protein arginine methyl transferase PRMT5.^17, 21–27^ Amongst these the combination of HDACs and histone methyltransferases provide the potential to convert the epigenetic state of target genes from an open to repressed chromatin state, with recruitment of G9A and EZH2 providing the capacity to mediate the establishment of repressive methylation marks at H3K9 and H3K27 residues.^17, 24, 28, 29^

In vitro models allow the sequential tracking of transitions between cell states during PC differentiation. In human models CD40L-based activation has provided a central platform for understanding PC differentiation.^30^ Here we explore the gene regulatory changes that characterize the transition between ABC and plasmablast states following removal from CD40L signaling in such a model.^5, 31^ We assess the relationship to genomic occupancy and epigenetic state linked to BLIMP1, IRF4 and XBP1 in plasmablasts. These data reveal a dichotomy in association of these core transcription factors to the pattern of gene expression change that characterizes the ABC to plasmablast transition.

## Materials & Methods

### Reagents

For the *in vitro* cell stimulation and maintenance reagents were as follows: human IL-2 (Roche); IL-21 (PeproTech). Goat anti-human F(ab’)_2_ fragments (anti-IgM & IgG) (Jackson ImmunoResearch); Lipid Mixture 1 chemically defined (200X) and MEM Amino Acids Solution (50X) (Sigma). For G9A inhibition: UNC0638 (Cayman Chemical). For cell proliferation: Carboxyfluorescein diacetate succinimidyl ester (CFSE) (Sigma).

### Donors and cell isolation

Peripheral blood was obtained from healthy donors after informed consent. Mononuclear cells were isolated by Lymphoprep (Axis Shield) density gradient centrifugation. Total B-cells were isolated by negative selection with the Memory B-cell Isolation Kit (Miltenyi Biotec). Memory-enriched B cells fractions were isolated by negative selection following incubation of total, negatively selected B cell fractions with CD23 Biotin and anti-Biotin Microbeads (Miltenyi Biotec)

### Cell cultures

24-well flat-bottom culture plates (Corning) and Iscove Modified Dulbecco Medium (IMDM) supplemented with Glutamax and 10% heat-inactivated fetal bovine serum (HIFBS, Invitrogen) were used. Day-0 to day-3 – B cells were cultured at 2.5 × 10^5^/ml with IL-2 (20 U/ml), IL-21 (50 ng/ml), F(ab’)_2_ goat anti-human IgM & IgG (10 µg/ml) on γ-irradiated CD40L expressing L cells (6.25 × 10^4^/well). Day-3 to day-6 – At day-3 cells were detached from the CD40L L-cell layer and reseeded at 1 × 10^5^/ml in media supplemented with IL-2 (20 U/ml), IL-21 (50 ng/ml), Hybridomax hybridoma growth supplement (11 µl/ml), Lipid Mixture 1, chemically defined and MEM Amino Acids solution (both at 1x final concentration). For UNC0638 experiments, ABCs were treated at day-3 with inhibitor at the indicated concentration (generally 2 μM), vehicle control (DMSO) or with standard conditions as indicated. Cells were sampled at the indicated time points without further addition of inhibitor.

### Flow cytometric analysis and microscopy

Cells were analysed using 4- to 6-color direct immunofluorescence staining on a BD LSR II flow cytometer (BD Biosciences). Antibodies used were: CD19 PE (LT19) and CD138 APC (B-B4; Miltenyi Biotec); CD23 APC (M-L233), CD27 FITC (M-T271), CD38 PE-Cy7 (HB7; BD Bioscience); CD20 efluor V450 (2H7) (eBioscience). Controls were isotype-matched mouse mAbs. Dead cells were excluded by staining with 7-AAD (BD Biosciences). Autophagy was detected with the Cyto-ID^®^Autophagy Detection Kit (Enzo Life Sciences). Absolute cell counts were performed with CountBright beads (Invitrogen). Cell populations were gated on FSC and SSC profiles for viable cells determined independently in preliminary and parallel experiments. Analysis was performed with BD FACSDiva Software 8.0 (BD Biosciences) and FlowJo v10 (FlowJo LLC).

### Gene expression analysis

RNA was extracted with TRIzol™ (Invitrogen) and amplified using Illumina® TotalPrepTM-96 RNA Amplification Kit (Life Technologies). Resultant cRNAs were hybridized onto HumanHT-12 v4 Expression BeadChips (Illumina) according to manufacturer’s instructions, scanned with the Illumina BeadScanner and initial data processing carried out using the Illumina GenomeStudio. For details of normalization and analysis please see Supplemental Methods.

### PGCNA and Signature enrichment analysis

See Supplemental Methods, for details of PGCNA, in brief informative genes are used to calculate Spearman’s rank correlations for all gene pairs. For each gene (row) in a correlation matrix only the 3 most correlated edges per gene are retained. The resulting matrix M is made symmetrical. The correlation matrices are clustered using a community detection algorithm and the best (judged by modularity score) used for downstream analysis.^32^ Gene signature analysis for modules was performed using a hypergeometric test against a curated sets of signatures.^33–37^

### Western blot, ChIP and ChIP-seq

At the indicated time points, primary cells were harvested, washed in PBS and lysed in Laemmli buffer to generated whole cell lysates. Western blots were performed using the following antibodies: Blimp-1 (R23),^38^ H3(ab1791), H3K9me2 (ab1220), H3K27me3 (07-449), and autophagy detection kit (Cell Signaling). ChIP was performed as described.^39^ Antibodies used were Blimp-1(R21),^38, 40^ IRF4 (sc - 28696X), XBP1 (Biolegend, 619502), CTCF(07-729), H3K4m3(04-745), H3K9me2(ab1220), H3K27me3(07-449), H3K27Ac(ab4729). ChIP-seq libraries were prepared using the MicroPlex Library Preparation™ kit (Diagenode) for ChIP, size selected using AMPure XP beads (Beckman Coulter) and run on an Illumina Hiseq 2500.

### ChIP-seq data analysis and motif detection

For more detail see Supplemental Methods. Trimmed reads were aligned with Bowtie2,^41^ and analysed for peaks using GEM and MACS2 with overlapping peak sets retained.^42, 43^ Peak overlaps were determined using a clustering approach such that any peak centre <500bp from an index peak centre were considered part of an overlapping cluster (See Supplemental Methods). *De novo* motif detection was performed with HOMER.^44^

The high-confidence peak sets for BLIMP1, IRF4 and XBP1 along with the Union set (overlap of individual high-confidence BLIMP1, IRF4 and XBP1 peaks) were analysed. Peaks were normalised and extended to their estimated fragment length. Scores per region were calculated for a ± 1000bp region. The resulting matrix was k-means clustered and visualised, see Supplemental Methods for details.

Data sets are available with GEO accession GSE142494.

### Ethical approval

Approval for this study was provided by UK National Research Ethics Service via the Leeds East Research Ethics Committee, approval reference: 07/Q1206/47.

## Results

### Activated B-cells encompass a growth state of B-cell differentiation

In order to explore the ABC to plasmablast transition we initially performed a time-course gene expression experiment from total human peripheral blood B-cells and explored the data with parsimonious gene correlation network analysis (PGCNA), an efficient computational approach recently developed in our lab.^5, 32^ Cells were sampled at day-0 (resting B-cell) on day-3 after activation with CD40L, anti-BCR and cytokines (ABC) and then at intervals of 3h, 6h, 12h, 24h, 48h and 72h (plasmablast) after transition into conditions (continued IL2 and IL21 only) that support the ABC/plasmablast transition. The network representing gene expression change over this differentiation window was comprised of 20 modules (Figure 1A, Supplemental Figure 1, Supplemental Table 1 and https://mcare.link/g9a). Gene signature and ontology enrichment was used to assess biological functions associated with each module which illustrated effective separation of known biological pathways between modules (Figure 1B, Supplemental Figure 2 and Supplemental Table 2). Summary designations for each module were derived from these enrichments. Overlaying the expression z-scores from the differentiation series then allowed visualization of the transitions in gene expression across the network as differentiation progressed (Figure 1C and D, and interactive network resources).

**Figure 1.**
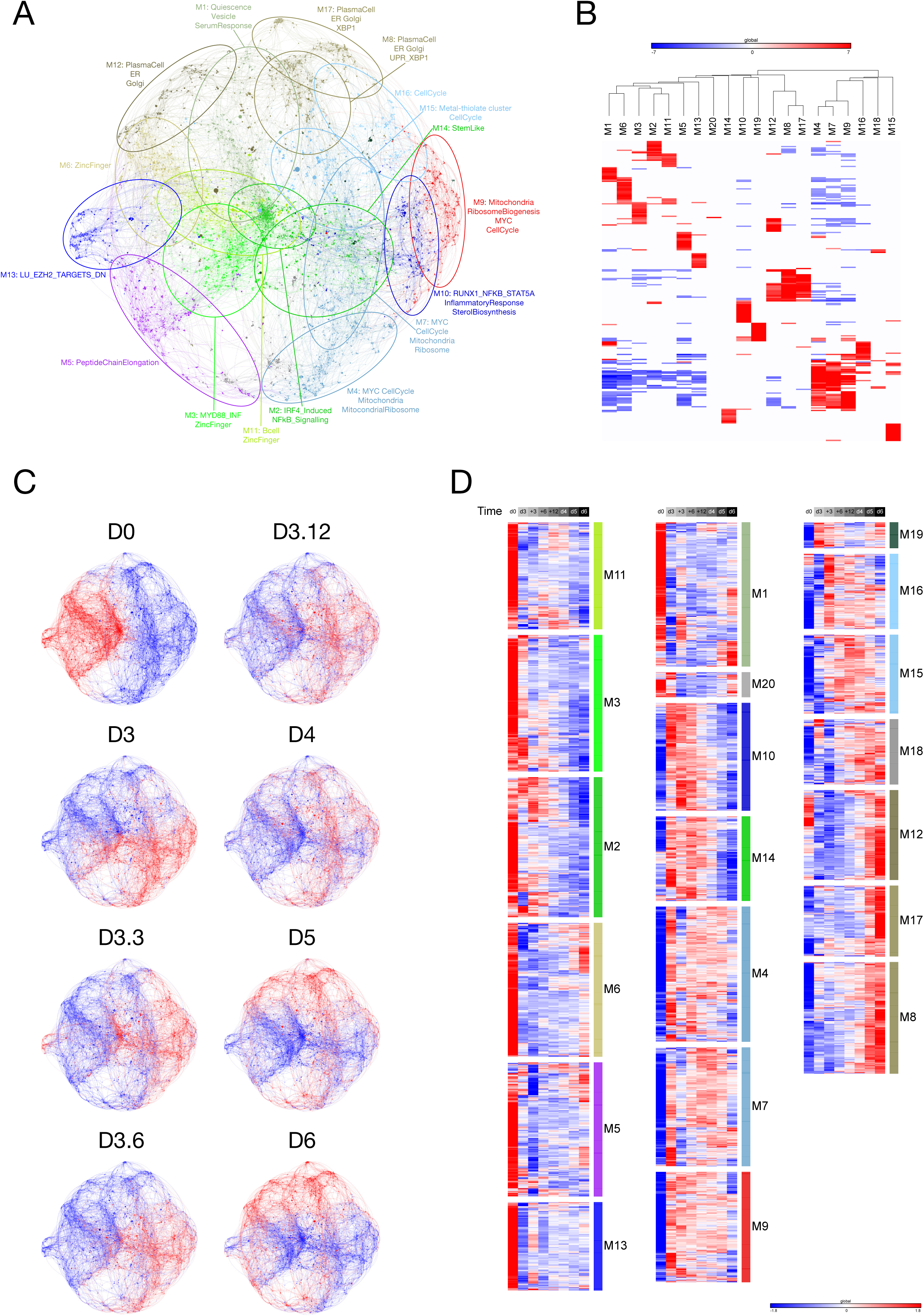
Application of PGCNA to time course gene expression data of total peripheral blood B-cell differentiation to PB state. **(A)** Network representation of the modular pattern of gene expression during the transition of B-cell to PB. Module designations derived from gene ontology enrichment indicated with color code and ovals. Interactive version (https://mcare.link/G9A_Paper_Total-Std-Mod) **(B)** Heatmap summary representation of gene ontology and signature separation between network modules (filtered FDR <0.05 and ≥ 5 and ≤ 1000 genes; selecting the top 15 most significant signatures per module). Significant enrichment or depletion illustrated on red/blue scale, x-axis (signatures) and y-axis (modules). Hierarchical clustering according to gene signature enrichment. For high-resolution version and extended data see SF2 and Supplemental Table 2. **(C)** Overlay of gene expression z-scores for all genes in the network shown in blue (low) to red (high) z-score color scale. Day 0 (D0) provides the starting reference point for the sequential expression patterns observed at the subsequent time points indicated following decimal point for samples between D3 and D4. For interactive versions of all networks go to https://mcare.link/g9a **(D)** Heat map displaying the pattern of gene expression across the time course module numbers indicated on the right, z-score gene expression blue (−1.8 low) red (+1.8 high) color scale as indicated right lower edge. Showing the median expression across 3 donors per timepoint. Modules divided into three broad categories of kinetics on at D0 going off, transient expression between D0 and D6, upregulated at late time points.

Considering the temporal transitions between the three primary cell states, resting B-cells preferentially express modules enriched in genes characteristic of the B-lineage and chromatin regulators (M6, M11), peptide chain elongation (M5), a subset of the secretory apparatus related to the Golgi and glycoprotein biosynthesis (M12) as well as genes linked to endosomal vesicles and quiescence signatures (M1). Modules shared between the resting B-cell and ABC state (M2 and M3) link features of the B-lineage to characteristic elements of the NFkB and cytokine response pathways. At the ABC state the modules expressed preferentially in resting B-cells were repressed, while elements of modules M2 and M3 are retained. This was accompanied by enhanced expression of modules on the one hand characteristic of the signaling pathway inputs (M10) and three modules of genes related to cell growth and division with a dominant impact of MYC and E2F target gene expression (M4, M7 and M9). The separation of these three modules sharing common linkage to MYC and E2F targets reflects relative enrichment of distinct biological processes such as hallmark signatures of cell cycle G2/M checkpoint (M4 and M7), DNA repair and mRNA related metabolic processes (M7), and non-coding RNA, rRNA and telomerase/Cajal body RNA localization (M9). Thus, the ABC state was characterized by a dominant signature of MYC and E2F related growth programs and sustained evidence of activating input signals.

By contrast the eventual plasmablast state saw a silencing of B-cell modules (M2, M3 and M11), signal input modules (M10 and components of M2) and cell growth related modules (M4, M7 and M9) of the ABC state. These changes are paralleled by enhanced expression of modules related to the UPR and XBP1 signaling (M8), and the ER including additional XBP1 targets (M17). Additionally, several modules expressed in the quiescent B-cell state are re-expressed in the plasmablast state including genes related to the Golgi apparatus (M12), chromatin regulators including demethylases (M6), and the peptide chain elongation enriched module (M5). Thus, dominant growth programs are contained in the ABC state, while plasmablast and resting B-cell share gene expression related to functional pathways.

### Transition from loss of signal input to cell cycle and secretory modules

Intermediate time-points provide a more detailed view of the transitional states between signaling and growth programs and secretory functional programs that characterize the ABC to plasmablast transition (Figure 1C and D). The most immediate gene expression changes reflect the removal of CD40L and BCR signals that accompany transfer into renewed cytokine conditions. Within 3h a down-modulation amongst specific signaling pathway genes was seen including *TNFAIP3* and *RGS1* residing primarily in module M2 which is shared between the resting B-cell and ABC state (Supplemental Figure 3). Genes in the signal response module M10 such as *BATF* showed a slightly more delayed repression but decayed rapidly from 6h onward and were essentially silenced by 24h (Supplemental Figure 3). These kinetics contrast with MYC and cell cycle related gene expression modules of the ABC state (M4 and M7) which were maintained to 48h. An additional module of cell cycle linked genes (M16) which is relatively weakly expressed in the ABC state and is particularly enriched for genes related to mitotic cell cycle, shows increased expression from 3-6h and includes both the proliferation related transcription factors *FOXM1* and *MYB* and the classical G1/S phase marker *MKI67* (Supplemental Figure 3). Consistent with the plasmablast population remaining in cell cycle the expression levels of such proliferation-associated genes, while no longer at peak levels, remained considerably higher than in the initial quiescent B-cell state. These patterns of gene expression are consistent with the overall pattern of cell division kinetics and population cell number of the differentiation series, which shows a modest number of divisions over the first three days of activation, followed by a very rapid proliferative phase after release from CD40L (Supplemental Figure 4). This has parallels with the pattern of a growth phase preceding rapid division in murine models of the germinal centre response.^45, 46^

Within the transitional window the central drivers of transcriptional reprogramming *IRF4, PRDM1* (encoding BLIMP1) and *XBP1* show similar patterns overall with some upregulation at day-3 relative to day-0 and subsequent substantial increase over the following 72h (Supplemental Figure 3). Differentiation and initial secretory pathway gene expression is enriched in module M8 which includes these key elements of the PC state along with UPR target genes which principally initiate expression from 12-24h onward. Secretory program gene expression extends into modules enriched for distinct secretory pathway elements (M17 and M12), which include genes whose eventual peak expression is characteristic of the mature quiescent PC state.^5^

### Dynamics of gene expression for memory B-cells at the ABC to plasmablast transition

Memory B-cells provide a source of plasmablasts capable of generating long-lived PCs in vitro.^31^ We therefore next considered differentiation of memory B-cells in isolation, starting from the ABC state and using the same gene expression time course approach. The focused memory B-cell network was comprised of 21 modules that showed significant enriched biology (Figure 2A and B, Supplemental Figure 5 and 6, Supplemental Table 3 and 4). These followed the three general patterns concordant with the total B-cell differentiation: early silencing, transient expression and late induction across the ABC to plasmablast transition (Figure 2 C and D and https://mcare.link/g9a). Modules reflecting the MYC regulated growth program (m.M3), ribosome subunits and peptide chain elongation (m.M4) and the activation signal response (m.M5 and m.M8) dominated the ABC state. These were followed by transient upregulation of cell cycle related modules (m.M7 and m.M9) including the genes *MYB, MKI67*, and *FOXM1*, through to the induction of secretory program components (m.M1 and m.M2). Across the time course the largest variance was seen in genes belonging to m.M1, m.M5 and m.M8. While some divergence in module composition was evident between the total and memory B-cell related gene expression patterns, the overall progression in gene expression was highly similar with a transient wave of proliferative gene expression following the decay of input signals, and onset of secretory program from 24-48h after release into transitional conditions.

**Figure 2.**
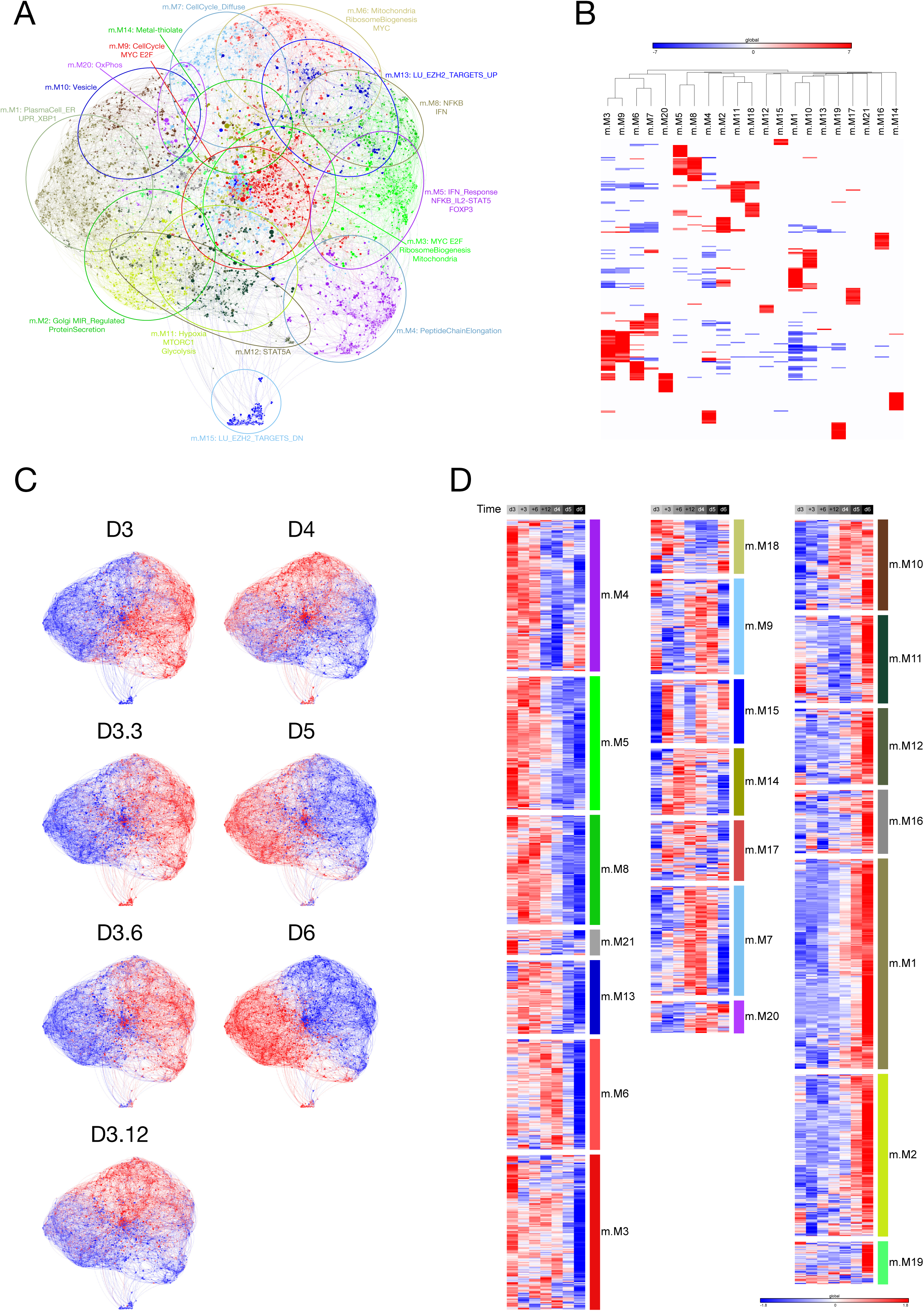
Application of PGCNA to time course gene expression data of memory B-cell differentiation from ABC to PB state. **(A)** Network representation of the modular pattern of gene expression during the transition of memory B-cell derived ABCs to PB state. Module designations derived from gene ontology enrichment indicated with color code and ovals. (Interactive version https://mcare.link/G9A_Paper_memB-Std-Mod) **(B)** Heatmap summary representation of gene ontology and signature separation between network modules (filtered FDR <0.05 and ≥ 5 and ≤ 1000 genes; selecting the top 15 most significant signatures per module). Significant enrichment or depletion illustrated on red/blue scale, x-axis (signatures) and y-axis (modules). Hierarchical clustering according to gene signature enrichment. For high-resolution version and extended data see SF2 and Supplemental Table 4. **(C)** Overlay of gene expression z-scores for all genes in the network shown in blue (low) to red (high) z-score color scale. Day 3 (D3) provides the starting reference point for the sequential expression patterns observed at the subsequent time points indicated following decimal point for samples between D3 and D4. For interactive versions of all networks go to https://mcare.link/g9a **(D)** Heat map displaying the pattern of gene expression across the time course module numbers indicated on the right, z-score gene expression blue (−1.8 low) red (+1.8 high) color scale as indicated right lower edge. Showing the median expression across 3 donors per timepoint. Modules divided into three broad categories of kinetics: (left) on at D3 going off, transient expression between D3 and D6, upregulated at D6.

### BLIMP1 and IRF4 occupancy in memory derived plasmablasts

To explore the relationship of TF occupancy patterns and gene regulation, we focused our analysis on the memory B-cell derived plasmablast population performing ChIP-seq for BLIMP1 and IRF4. We identified 4244 BLIMP1 occupancy sites of which the majority (69%) fell within intronic and intergenic regions and close to a quarter in promoter regions. For IRF4 we identified 9259 peaks of which a greater proportion (44%) fell within promoter regions and just under half in inter- or intra-genic regions (Figure 3A and Supplemental Table 5). Thus, IRF4 displayed a greater tendency for promoter occupancy. Of the total peak set 1807 regions were co-occupied by both factors (Figure 3B).

**Figure 3.**
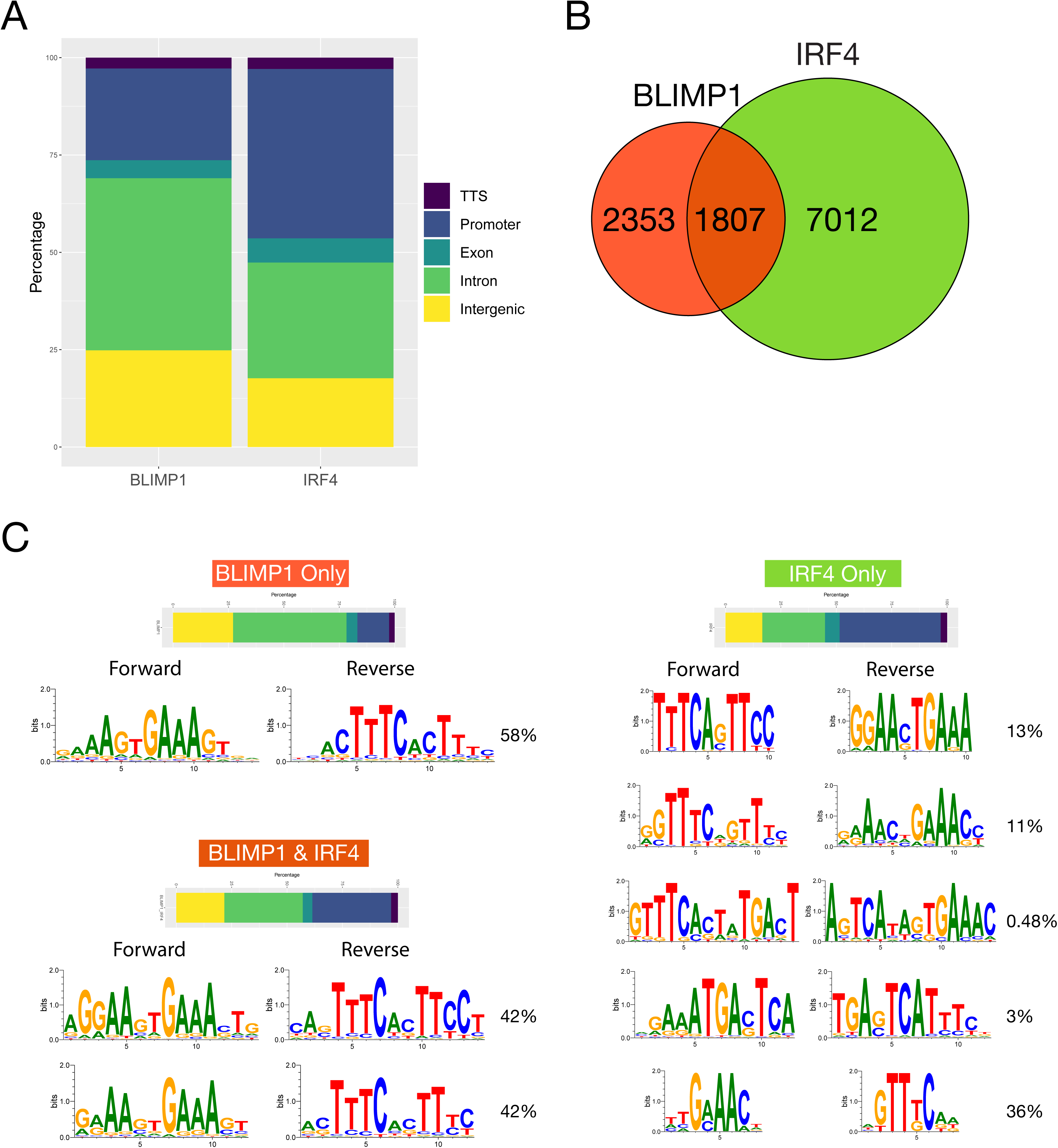
IRF4 and BLIMP1 occupancy in human PBs. **(A)** Relative distribution of BLIMP1 (left) and IRF4 (right) peaks identified in human memory B-cell derived PBs divided according to genomic distribution, transcription termination site (TTS), promoter, exonic, intronic and intergenic as indicated in the color code to the right of the stacked bar graph. **(B)** Venn diagram depiction of BLIMP1 and IRF4 binding site overlap genome wide. **(C)** Relative genomic distribution and de novo motifs discovered at sites of BLIMP1-only, IRF4-only and BLIMP1/IRF4 overlapping occupancy. Shown are the genomic distribution as stacked bar graph color-coded as in (A) and the most significantly enriched motifs with percentage of peak regions with match to represented motif variant to the right. For IRF4-only motifs the frequencies of variant IRF4 motif types are shown from top to bottom ISRE, EICE, AICEv1, AICEv2 and minimal IRF consensus.

De novo motif detection of occupied sites, confirmed the dominant enrichment of motifs matching the established consensus sequence (Figure 3C). The BLIMP1 and the IRF family share an evolutionarily conserved partial overlap in motif preference. We and others have previously shown that BLIMP1 and IRFs differ in their preference for the first position of the GTG triplet in the shared consensus, AA<u>G</u>TGAAA<u>G</u>T, where selection of a C rather than G disfavors BLIMP1 occupancy.^17, 47, 48^ This observation was supported in de novo motif analysis of individual or co-occupied sites. Furthermore, across IRF4 occupied sites, those occupied with BLIMP1 selected more strongly for ISRE and EICE consensus sequences, while a greater diversity of IRF4 binding motifs including AICE variant 1 and 2 were recovered at sites occupied by IRF4 alone. Notably at these regions the CTCF/BORIS motif was also significantly enriched (Supplemental Figure 7A). BLIMP1 and IRF4 therefore showed partially overlapping genomic occupancy in human plasmablasts, selecting for a subset of closely related binding motifs, with independent occupancy by each factor enriching for preferred variations. These selections were also linked to significant differences in co-occurring secondary motifs with BORIS/CTCF motifs preferentially linked to IRF4 occupancy in the absence of BLIMP1.

### BLIMP1, IRF4 and XBP1 occupy distinct regulatory element clusters

To profile the epigenetic state associated with IRF4 and BLIMP1 occupancy and relate this to the additional elements of the transcriptional network controlling PC differentiation we performed ChIP-seq for H3K4me3, H3K27ac, H3K9me2, CTCF and XBP1. For both CTCF and XBP1 we recovered the appropriate primary TF motif (Supplemental Figure 7B and C). XBP1 ChIP-seq provided the most limited peak set but with the established XBP1 consensus G(C/A)CACGT as the most significantly enriched motif. At a subset of sites a CCAAT box was also evident (Supplemental Figure 7C) and together these comprise the composite ER Stress Response Element (ERSE).^49^

We then used the union of genomic regions that were occupied by XBP1, IRF4 or BLIMP1 to assess recurring patterns of chromatin marks and CTCF occupancy (Figure 4). We used K-means to resolve 6 regulatory clusters amongst these peak regions. This encompassed three regulatory clusters of highly active regions with either 5’ or 3’ skewing of histone marks, or symmetric distribution around the TF site (U.K1-3), consistent with the general observation of widespread heterogeneity of histone modifications around TF sites.^50^ These clusters were promoter biased and relatively enriched for IRF4 and XBP1 occupancy. The fourth regulatory cluster was distinctively associated with CTCF occupancy and enriched for IRF4 binding relative to BLIMP1 or XBP1 (U.K4). The remaining two regulatory clusters were linked to weak (U.K5) or absent (U.K6) active histone marks and a broad symmetric distribution of the repressive chromatin marks around the peak centre. BLIMP1 binding was relatively enriched at both these clusters but in particular for cluster U.K6 which was also relatively depleted of XBP1 and CTCF binding.

**Figure 4.**
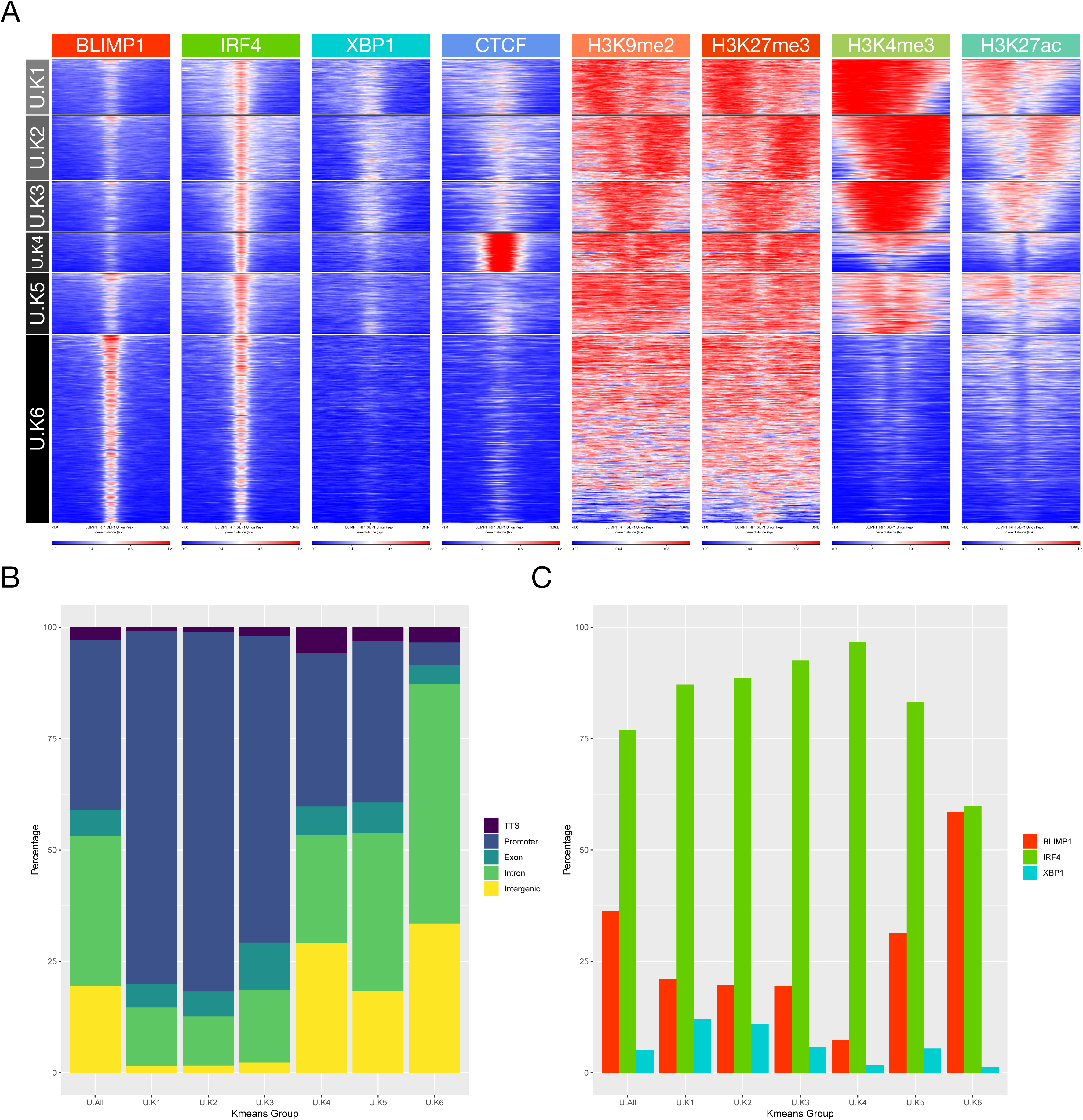
Epigenetic patterns associate with core TF occupancy at the PB state. **(A)** Deep tools heatmap representation of K-means clustered integrated ChIP-seq data from the PB state. Data is clustered across the union of peaks for IRF4, BLIMP1 and XBP1 and encompassing data for CTCF, H3K9me2, H3K27me3, H3K4me3 and H3K27ac from equivalent cells. 6 regulatory clusters are derived designated K1-K6 on the left. **(B)** Relative distribution of K-means clusters K1-K6 derived from (A) according to genomic distribution, transcription termination site (TTS), promoter, exonic, intronic and intergenic as indicated in the color code to the right of the stacked bar graph. **(C)** Percentage occupancy of individual TF binding across the K-means clusters derived (A) for each of the TFs indicated by the color code to the right of the figure (BLIMP1-red, IRF4-green, XBP1-blue).

Repeating the analysis centering on occupancy by each TF independently reinforced these differential patterns (Supplemental Figure 8, 9 and 10). XBP1 associated primarily with open active chromatin in promoter regions in the absence of BLIMP1. By contrast BLIMP1 was preferentially enriched in relatively inactive/repressed chromatin fractions comprising close to 70% of peak regions, either alone or with IRF4 co-occupancy. Twenty percent of BLIMP1 binding was linked to active chromatin in promoter biased regions. IRF4 bridges these patterns binding primarily at active regions with promoter enrichment, as well as in relatively silent chromatin enriched for intergenic and intronic regions and BLIMP1 occupancy. IRF4 in the absence of BLIMP1 or XBP1 showed a distinct association with CTCF, confirming the results of *de novo* motif analysis, and contrasting with XBP1 and BLIMP1 which associated infrequently with CTCF. Thus, each of the core TFs of the plasmablast state is linked at genome-wide level to a distinct pattern of epigenetic co-association.

### Modular gene expression link to TF occupancy and regulatory element clusters

To gain further insight into the link between regulatory clusters and expression patterns, we tested the association of the six regulatory clusters derived from the integrated and transcription factor specific analyses against the patterns of gene regulation observed in the memory B-cell PGCNA network. To do this we focused on occupancy proximal to a gene promoter (10kb no intervening TSS) as indicative of a potential regulatory interaction, looking for enrichment of such events across all genes in a module relative to background of all genes in the network. These analyses demonstrated that the regulatory element clusters defined using either integrated data or specifically for each transcription factor were significantly and differentially linked to the modular patterns of gene expression defined by PGCNA, in a fashion which was concordant with both known transcription factor biology and associated epigenetic state (Figure 5). Regulatory element clusters are significantly linked to the most characteristic and variant modules of gene expression across the network including both induced and repressed genes. However, it is also evident that several gene expression modules that characterize the ABC/plasmablast transition lack positive association with the regulatory element clusters linked to IRF4, BLIMP1, or XBP1. This supports the likely existence of additional yet to be defined transcriptional regulators active during this differentiation window, and responsible for the control of such modules.

**Figure 5.**
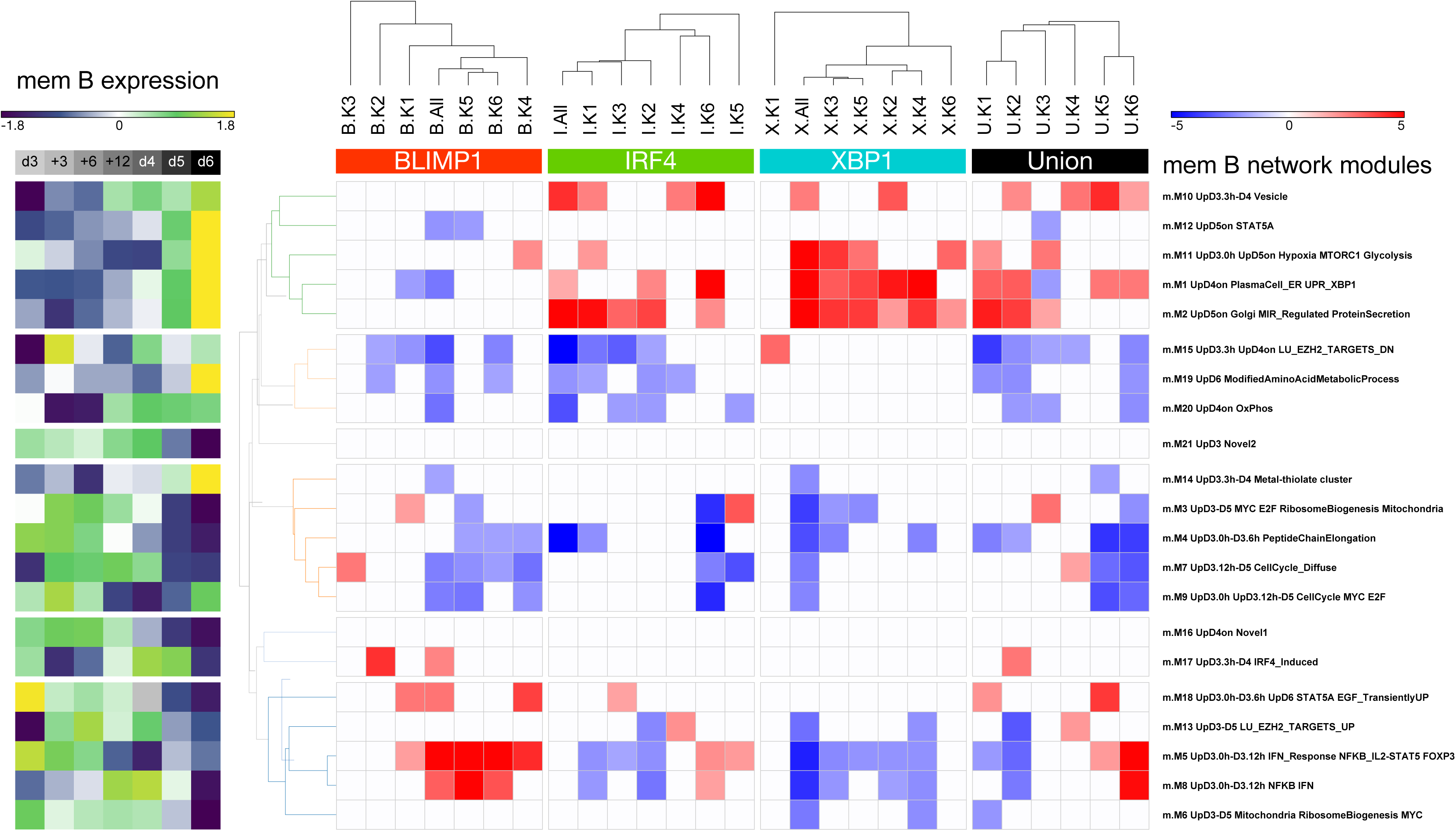
Integration of gene regulatory modules of the ABC to PB transition with TF occupancy patterns and epigenetic state. Shown is the signature enrichment heatmap displaying the enrichment/depletion of the memory B-cell PGCNA expression modules (Figure 2) for the genes associated with the TF peaks in the K-means clusters of epigenetic state (Figure 4). The significance of association between TF occupancy and genes belonging to a network module is shown as a z-score color scale (−5:blue to +5:red) divided across the top according to hierarchical clustering of K-means modules (z-scores with a p-value > 0.05 were set to 0). The results are shown in the order BLIMP1 K-means clusters (B.All & B.K1-6), IRF4 K-means clusters (I.All & I.K1-6), XBP1 K-means clusters (X.All & X.K1-6), Union K-means clusters (U.All & U.K1-6). On the right side the module identity is indicated. On the left side of the figure the median expression pattern of the module is shown across the time course as a z-score (−1.8:dark-blue to 1.8:yellow). For signature enrichment analysis the TF peaks within each k-mean set were associated with the nearest gene (≤ 10kb), and these genes were used for a hypergeometric test against the PGCNA module gene-sets.

Considered from the point of view of gene expression, modules that are induced at the plasmablast stage and encompass core phenotypic and functional pathways of this state (m.M1, m.M2, m.M10, and m.M11) link to regulatory element clusters characterized by active marks, promoter enrichment and IRF4 and/or XBP1 occupancy. These modules are neutral or anti-correlated with BLIMP1 occupancy, with the exception of the m.M11 module linking to B.K4 (symmetric active) with weak significance. Reciprocally modules characteristic of ABCs and repressed at the plasmablast state (m.M5, and m.M8) are correlated with regulatory element clusters linked to BLIMP1 occupancy, and inactive/repressive chromatin states. Such modules of gene expression are reciprocally either neutral or anti-correlated with respect to the active regulatory clusters. Thus, the regulatory element clusters defined by IRF4, XBP1 and BLIMP1 occupancy provide a coherent picture and link in a dichotomous fashion to the key elements of the gene expression network of the ABC to plasmablast transition.

### G9A inhibitor UNC0638 impacts on the efficiency of ASC generation

BLIMP1 can recruit EHMT2/G9A, the H3K9 directed methyltransferase which catalyzes the H3K9me2 modification. The association between BLIMP1 occupancy and the regulatory cluster (U.K6) characterized by relative enrichment of H3K9me2 modification pointed to a potential role for this modification in the repression of genes linked to the ABC state. We therefore aimed to explore the relative contribution of H3K9me2 modification taking advantage of the selective pharmacological inhibitor of G9A, UNC0638.

We initially evaluated the impact of UNC0638 treatment on the functional characteristics of the ABC to plasmablast transition. We identified a dose of UNC0638 that was sufficient to impair features of phenotypic differentiation and to impact on global H3K9me2 levels across the 72h of culture (Supplemental Figure 11A). G9A inhibitors have been reported to induce autophagy, this was also observed in the B-cell response to this G9A inhibitor with rapid induction of features consistent with autophagy in ABCs following UNC0638 treatment (Supplemental Figure 11B and C). Autophagy can have a protective role in PC differentiation,^51, 52^ and despite the stress response phenotypic differentiation showed limited differences with modest impairment in the downregulation of CD20 and the upregulation of CD38 (Supplemental Figure 11D). Furthermore, consistent with a delayed impact as would be anticipated with an epigenetic mechanism cell division was modestly impaired at day 4 (24h of treatment) and delayed by one generation at day 5 and 6 where the population mode for UNC0638 treated cells fell at generation 6 rather than 7 (Supplemental Figure 11E). Notably functional differentiation was largely maintained, with day 6 cells seeded at equivalent densities showing equal numbers of ASCs (Supplemental Figure 11F). Thus, at a dose of G9A inhibitor that reduced global methylation of H3K9me2 and induced the characteristic autophagic stress response, functional ASC differentiation remained largely intact providing a suitable condition in which to assess consequences of acute G9A inhibition.

### A subset of BLIMP1 bound genes linked to the ABC state is responsive to UNC0638

Despite the rapid induction of cellular stress response, there were no significantly differentially expressed genes following UNC0638 treatment until 24h after treatment, and substantial numbers of differentially expressed genes did not appear until 72h in day-6 plasmablast (Figure 6A, Supplemental Table 6). The impact of UNC0638 on autophagic stress response, which occurred early, is therefore not explained in this system by differential gene regulation. Furthermore, the delayed impact on gene expression is consistent with an epigenetic effect in this rapidly dividing population. We therefore tested BLIMP1 occupancy and associated epigenetic marks at the day-6 time point of maximal difference in gene expression.

**Figure 6.**
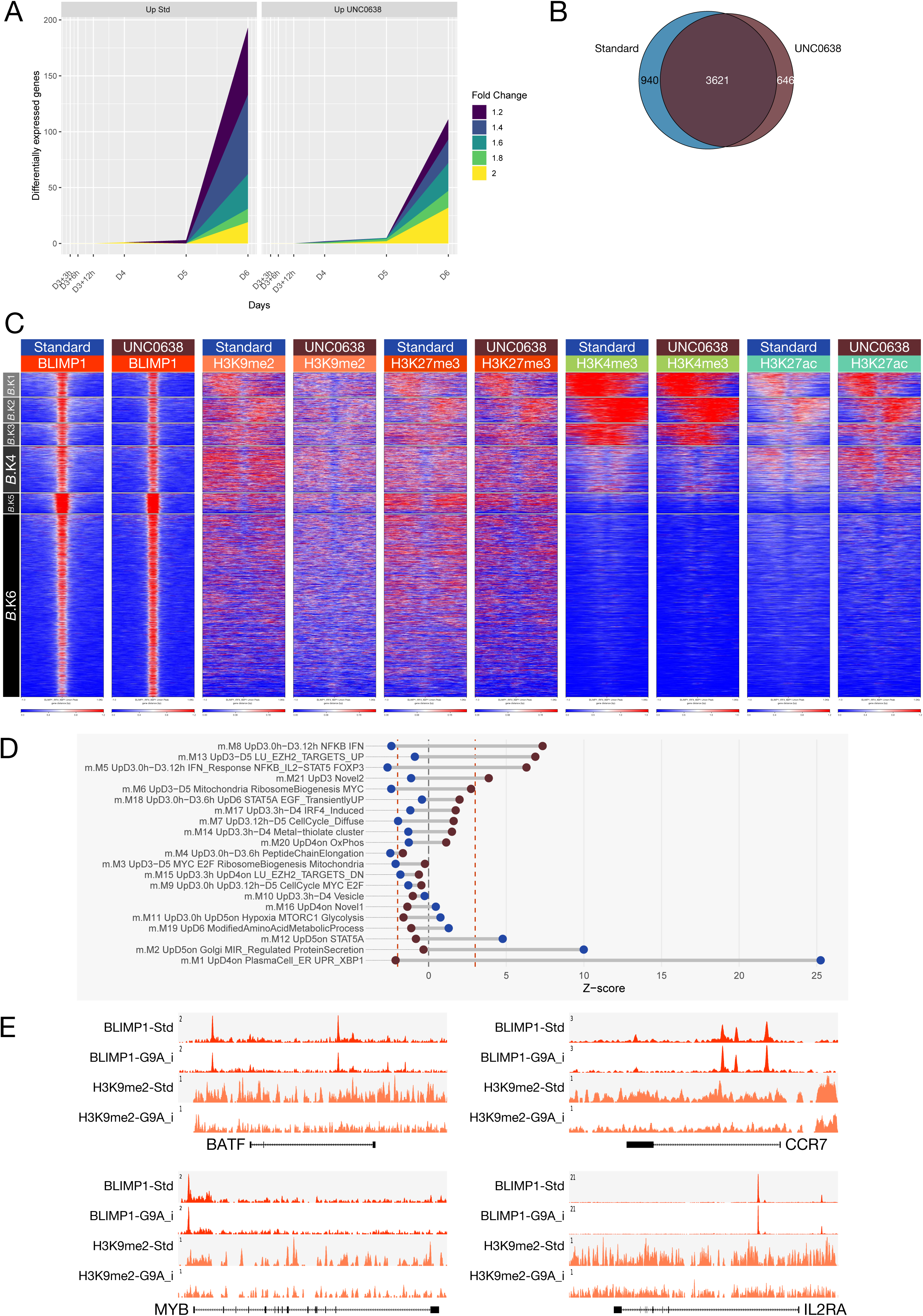
G9A inhibition with UNC0638 produces a focused impact on gene expression during PB differentiation. **(A)** Graphical representation of differential gene expression across a range of fold-change thresholds from 1.2+ to 2.0+ across a time course following UNC0638 treatment at D3 of memory B-cell differentiation. Number of differentially expressed genes across each fold-change threshold indicated by the color coded graphical representation, according to color code on the right of the figure. (y-axis number of significantly differentially expressed genes, x-axis time point in hours and days). Left graph genes upregulated in the absence of inhibitor, right graph genes upregulated in presence of inhibitor. Data derived from three independent donors. **(B)** The overlap of BLIMP1 occupancy in memory derived PBs in the absence (blue) or presence (brown) of UNC0638 treatment. **(C)** Deeptools heatmaps of K-means clusters derived from the union of BLIMP1 binding sites for standard and UNC0638 conditions and considering associated epigenetic marks as indicated for H3K9me2, H3K27me3, H3K4me3, and H3K27ac. **(D)** Dumbbell graph of the relative enrichment or depletion of differentially expressed genes shown in (A) against the modules of gene expression derived from the memory B-cell PGCNA network in Figure 2. Y-axis shows the order of modules ranked from most significantly enriched for genes upregulated in presence of UNC0638 through to most significantly enriched in the standard conditions. For each module the enrichments or depletion are shown for the genes upregulated in the presence of UNC0638 (brown circles) and genes upregulated in standard conditions (blue circles). These are plotted against the x-axis displaying Z-score of enrichment/depletion with the vertical dotted red lines indicating the point of FDR corrected significance (p-value<0.05). **(E)** Representative tracks for BLIMP1 and H3K9me2 ChIP-seq in standard and UNC0638 treated samples, as indicated for representative genes whose expression is increased in the presence of UNC0638.

Consistent with the maintenance of functional ASC differentiation, BLIMP1 occupancy was globally unimpaired in the presence of UNC0638 treatment (Figure 6B). Repeating the K-means clustering of all BLIMP1 peak regions (union of control/inhibitor-treated peaks), illustrated that overall patterns of active and repressed regulatory regions were maintained. Consistent with an on-target inhibitor effect there was a general reduction in H3K9me2 signal around, but not restricted to, BLIMP1 target sites (Figure 6C). Importantly H3K27me3, H3K4me3 and H3K27ac signals were not reduced in the presence of inhibitor. Thus, a global but partial impact of G9A inhibitor treatment on H3K9me2 state contrasted with a highly selective impact on differential gene expression.

We next mapped the gene expression changes onto the PGCNA network for the ABC to plasmablast transition. This demonstrated a focused effect for both genes differentially upregulated and downregulated by UNC0638 (Figure 6D). Genes that were expressed at higher levels in control conditions at day 6 were significantly enriched amongst genes in modules upregulated at the plasmablast stage m.M1, m.M2 and m.M12 and were most strongly enriched amongst genes in module m.M1 encompassing genes characteristic of the PC state. By contrast those genes expressed at higher level in the presence of UNC0638 (downregulated in standard conditions) were significantly skewed to modules with opposite kinetics during normal differentiation, that are repressed during plasmablast differentiation m.M5, m.M8 and m.M13 and that link to regulatory element clusters associated with BLIMP1 occupancy (m.M5 and m.M8), and relatively silenced chromatin state. Hence the impact of G9A inhibition on expression is concordant with the regulatory element clusters and consistent with a focused delay in differentiation related gene expression.

While UNC0638 mediated G9A inhibition is not selective for the nature of G9A targeting to chromatin, the pattern of gene expression changes supports a functional link with BLIMP1. Integrating differential expression with occupancy showed a significant skewing toward local BLIMP1 occupancy amongst genes upregulated in the presence of UNC0638 (2-fold-up: 14/32, 44% BLIMP1 bound vs 2-fold-down: 3/19, 16% BLIMP1 bound). The genes differentially upregulated included *IL2RA*, which has been previously defined as a BLIMP1 and G9A target in T-cells.^53^ Additionally, the gene set included characteristic transcription factors of the ABC state *BATF, MYB* and *RUNX3* as well as typical features of the ABC state *CCL22, CCR7* and *ARHGAP17* (Figure 6F).^54^ This coherent impact of G9A inhibition on the differentiation program provides evidence for a selective dependency: repression of components of the activated B-cell program that are also associated with local BLIMP1 occupancy are sensitive to G9A inhibition.

## Discussion

In the direct differentiation of B-cell to the PC state the two primary transitional states that can be defined are the activated B-cell and the plasmablast. Both of these are ephemeral cell states, the former linked to the extracellular cues driving the activation process, and the latter representing the penultimate stage poised for decision between cell death or entry into cell cycle quiescence and the completion of the PC differentiation program. This transition encompasses the cusp between the lymphoid and the antibody secreting states of the B-cell life cycle. Given the prevailing model of a broadly epistatic relationship between IRF4, BLIMP1 and XBP1 during the establishment of PC differentiation,^1^ we sought to explore how the reorganizing gene expression network in human B-cells related to the pattern of TF occupancy at the plasmablast state. Our data support the conclusion that these three TFs have quite distinct associations with both epigenetic state, and gene regulation. We have recently established an expression networking tool, PGCNA, which can be effectively applied to time course data sets.^5, 32^ Here, this approach allows the detailed definition of modules of gene co-expression accompanying the differentiation of either total peripheral blood B-cells or isolated memory B-cells between the ABC and plasmablast states. The integration of these expression modules with global occupancy patterns and associated epigenetic patterns supports the general segregation between IRF4 and XBP1 as activators of gene expression and BLIMP1 as a repressor during this transition. The key modules that define the PC state are statistically significantly enriched for association with local occupancy by IRF4 and XBP1 in association with active chromatin patterns, while BLIMP1 occupancy at the plasmablast stage is significantly associated with genes that are repressed in the ABC/plasmablast transition and overall BLIMP1 occupancy is anti-correlated with the key module characteristic of the plasmablast/PC state (e.g. m.M1). In the differentiation of murine PCs an expanded role for BLIMP1 in the direct control of secretory pathway components has been described.^13, 17^ This contrasts with the lack of significant enrichment for local BLIMP1 occupancy either with the most characteristics expression module of the plasmablast/PC state or modules encompassing the wider secretory pathway in human plasmablasts. However, we note that at an individual gene level BLIMP1 does occupy regulatory elements of select secretory pathway genes in human plasmablasts. For example, of the 45 secretory pathway genes linked to BLIMP1 occupancy in murine PCs, ^13^ seven (*SIL1, DPAGT1, ERP44, HSP90B1, PDIA3, PPP1R15A, SLC33A1*) showed evidence of BLIMP1 binding in human plasmablasts at the level of peak calling stringency used in our analysis (identifying 4244 BLIMP1 peaks). However, placing this in context, of the more discrete set of XBP1 targets identified (789 peaks) 24/45 of this list of secretory pathway components were occupied by XBP1. Thus, our data show that in human plasmablasts while a few secretory pathway genes have evidence of local BLIMP1 occupancy, this is not a statistically enriched global feature of the BLIMP1 occupancy pattern. In human plasmablasts the strongest associations for BLIMP1 are with occupancy at regulatory elements linked to predominantly repressive chromatin marks, and genes expressed at the ABC stage that are repressed in plasmablasts. These include key transcriptional regulators such as *SPIB* and *IRF8*,^55–57^ which are also observed as targets in murine cells and aligns with features of human B-cell lymphomas with BLIMP1 inactivation.^58, 59^

Secretory pathway genes in human plasmablasts show the most significant association with XBP1 occupancy. Our understanding of the importance of XBP1 to the establishment of the PC fate has shifted from central integration of the ER stress response via XBP1 with the differentiation process,^60, 61^ toward a more specialized role for XBP1 in optimization of PC secretory functions.^14, 16, 62^ Our data demonstrate that in human plasmablasts XBP1 selectively binds to active promoters of many secretory pathway genes. The kinetics of expression of these genes is in parallel with, rather than subsequent to, that of other characteristic phenotypic components of the PC state. Moreover, the module of gene expression with which XBP1 occupancy associates, initiates expression from 24h after release from CD40 signal and is highly enriched for previously established signatures of the UPR and ER-stress response. While an additional wave of UPR/ER-stress responsive gene regulation occurs as human plasmablasts complete the differentiation to the quiescent PC state,^5^ the data present here using a spliced XBP1 specific antibody point to a significant contribution for XBP1 activation via the classical IRE1 processing pathway at the initial point of secretory pathway commitment. Murine models of conditional XBP1 deletion have demonstrated that a contribution from XBP1 is not an absolute pre-requisite for the generation of phenotypic PCs but is required for establishment of optimal secretory function. These results may in part be explained by functional redundancy since additional TFs linked to the ER stress response are regulated during PC differentiation such as ATF6 and CREB3L2.^63, 64^

A notable finding in relation to IRF4 occupancy in human plasmablasts is a significant association with CTCF binding, which is substantially different in proportion for that observed for either BLIMP1 or XBP1. Indeed, co-occupancy by IRF4 and CTCF is linked to motifs that disfavor BLIMP1 binding. Recently an association between IRF4 and CTCF has been identified in Th17 cells at sites co-occupied with STAT3, BATF and BRD2.^65^ Of note in plasmablasts, at CTCF/IRF4 co-occupied sites, EICE motifs predominate which suggests a divergence from the binding mode in the Th17 context. It will be intriguing to explore whether the link between IRF4 and CTCF reflects a particular contribution to long range chromatin interactions in plasmablasts.

In order to regulate gene expression, BLIMP1 recruits epigenetic coregulators.^17, 21–27^ While an important role for EZH2 has been identified,^17, 25^ the relative contribution of G9A and in particular G9A in human cells has not been extensively studied.^66^ Small molecule inhibitors of epigenetic regulators provide tool compounds for exploring functional biology.^67^ Selective inhibitors of G9A have been developed and validated across a range of different cell types in particular in cancer cell lines. UNC0638 acts through competitive binding in the peptide substrate recognition pocket, while not competing with the cofactor s-adenosyl-methionine.^68^ A feature of inhibition of epigenetic repressors including G9A has been evidence that acute loss of function leads to selective impact on target gene regulation without globally impairing epigenetic programs.^69, 70^ G9A inhibition did not prevent the execution of the majority of the plasmablast program but led to a focused impact on gene expression consistent with a partial delay in differentiation. This included the gradual de-repression of genes which belonged predominantly within modules linked to BLIMP1 occupancy and silent epigenetic state. Our results thus support a particular role for G9A in silencing of components of the activation program. Notable amongst the specific de-repressed genes are the cytokine receptor gene IL2RA, which has previously been identified as a BLIMP1 and G9A targets in T-cells,^53^ and the transcription factor BATF. The latter is one of the more distinctive transcriptional regulators of the ABC state and provides an important co-factor for IRF4 and potentially IRF8 in this context.^59^

In summary this integrated analysis illustrates the connection between reorganizing gene expression and epigenetic states of the ABC to plasmablast transition in human B-cells, and reinforces the integration and functional segregation between IRF4, BLIMP1 and XBP1 during this process. The limited impact of acute global inhibition of an epigenetic co-regulator in this system underlines the robustness of the system driving differentiation of the PC state, while pointing to a specific role for G9A in suppression of components of the B-cell activation program.

## Acknowledgements

This work was supported by Cancer Research UK program grant (C7845/A17723), and a studentship of the Sultanate of Oman (to MAM).

We thank Ulf Klein for critical review of the manuscript.

## Supplemental Figure Legends

**Supplemental Figure 1. Accompanies Figure 1.** High resolution version of Figure 1A, network representation showing color coded modules and genes. Refer also to https://mcare.link/g9a for an interactive searchable version including Z-score expression overlays as in Figure 1C. For individual gene expression values and module gene lists see Supplemental Table 1.

**Supplemental Figure 2. Accompanies Figure 1.** High resolution version of Figure 1B. Heatmap of gene ontology and signature term enrichments linked to the PGCNA modules of the time course network analysis (filtered FDR <0.05 and ≥ 5 and ≤ 1000 genes; selecting the top 15 most significant signatures per module). For full signature enrichment lists, please see Supplemental Table 2. Modules are shown along the x-axis, and selected signature terms along the y-axis. Signature terms and modules are hierarchically clustered to illustrate relationships. Enrichment (red) and depletion (blue) of signatures are shown on color scale of z-score.

**Supplemental Figure 3. Accompanies Figure 1.** Violin plots of individual gene expression for selected genes showing expression Z-score on the y-axis and time point in days and hours along the x-axis for the indicated genes above each graph. Violin plots display the distribution (n=3 donors) along with median (blue square) and the IQR.

**Supplemental Figure 4. Accompanies Figure 1.** Proliferation and matching phenotypic maturation during resting B cells to PB transition. Left panels histograms flow plots of CFSE dilution time course (left panel) with matching density plots for CD19 vs CD20 and CD38 vs CD138 (middle and right panel). Numbers indicate percentages of total cell populations following single cell and viability discrimination with gates established using matching isotype controls. Cells were labelled at Day 0 and proliferation and phenotypical maturation was assessed at Day 0, Day 3 and then monitored every 24 hrs up to Day 6 in Std treatment conditions. Data representative of three biological replicates.

**Supplemental Figure 5. Accompanies Fig 2.** High resolution version of Figure 2A, network representation showing color coded modules and genes. Refer also to https://mcare.link/g9a for an interactive searchable version including Z-score expression overlays as in Figure 2C. For individual gene expression values and module gene lists see Supplemental Table 3.

**Supplemental Figure 6. Accompanies Figure 2.** High resolution version of Figure 2B. Heatmap of gene ontology and signature term enrichments linked to the PGCNA modules of the time course network analysis of memory B-cell differentiation (filtered FDR <0.05 and ≥ 5 and ≤ 1000 genes; selecting the top 15 most significant signatures per module). For full signature enrichment lists, please see Supplemental Table 4. Modules are shown along the x-axis, and selected signature terms along the y-axis. Signature terms and modules are hierarchically clustered to illustrate relationships. Enrichment (red) and depletion (blue) of signatures are shown on color scale of z-score.

**Supplemental Figure 7. Accompanies Figure 3. (A)** de novo motif data for BLIMP1 only, BLIMP1_IRF4 and IRF4_only peak regions showing most significantly enriched motifs returned by HOMER analysis. **(B)** de novo motif analysis of CTCF peak set **(C)** de novo motif analysis of XBP1 peak set. For each data-set shows the top 5 primary HOMER de novo motifs. Above each motif set is a stacked bar chart showing peak genomic localization percentages for: transcription termination site (TTS), promoter, exonic, intronic and intergenic; as indicated in the colour code in the bottom right of the figure.

**Supplemental Figure 8. Accompanies Figure 4. (A)** Deep tools heatmap representation of K-means clustered integrated ChIP-seq data from the PB state. Data is clustered across the set of BLIMP1 peaks and encompassing data for CTCF, H3K9me2, H3K27me3, H3K4me3 and H3K27ac from equivalent cells. 6 regulatory clusters are derived designated K1-K6 on the left. **(B)** Relative distribution of BLIMP1 K-means clusters K1-K6 derived from (A) according to genomic distribution, transcription termination site (TTS), promoter, exonic, intronic and intergenic as indicated in the color code to the right of the bargraph. **(C)** Percentage occupancy of individual TF binding across the K-means clusters derived (A) for each of the other TFs indicated by the color code to the right of the figure (IRF4-green, XBP1-blue).

**Supplemental Figure 9. Accompanies Figure 4. (A)** Deep tools heatmap representation of K-means clustered integrated ChIP-seq data from the PB state. Data is clustered across the set of IRF4 peaks and encompassing data for CTCF, H3K9me2, H3K27me3, H3K4me3 and H3K27ac from equivalent cells. 6 regulatory clusters are derived designated K1-K6 on the left. **(B)** Relative distribution of IRF4 K-means clusters K1-K6 derived from (A) according to genomic distribution, transcription termination site (TTS), promoter, exonic, intronic and intergenic as indicated in the color code to the right of the bargraph. **(C)** Percentage occupancy of individual TF binding across the K-means clusters derived (A) for each of the other TFs indicated by the color code to the right of the figure (BLIMP1-red, XBP1-blue).

**Supplemental Figure 10. Accompanies Figure 4. (A)** Deep tools heatmap representation of K-means clustered integrated ChIP-seq data from the PB state. Data is clustered across the set of XBP1 peaks and encompassing data for CTCF, H3K9me2, H3K27me3, H3K4me3 and H3K27ac from equivalent cells. 6 regulatory clusters are derived designated K1-K6 on the left. **(B)** Relative distribution of XBP1 K-means clusters K1-K6 derived from (A) according to genomic distribution, transcription termination site (TTS), promoter, exonic, intronic and intergenic as indicated in the color code to the right of the bargraph. **(C)** Percentage occupancy of individual TF binding across the K-means clusters derived from (A) for each of the other TFs indicated by the color code to the right of the figure (BLIMP1-red, IRF4-green).

**Supplemental Figure 11. Accompanies Figure 6. Functional analysis of G9A inhibition with UNC0638 (A)** WB of total H3K9me2 in activated B-cells differentiating under standard conditions (-), with vehicle control DMSO, or after treatment at day-3 with UNC0638 at 2 μM concentration. Cells were sampled at the indicated time points. Upper blot H3K9me2, lower blot total H3 loading control. **(B)** WB of LC3A/B-I and LC3A/B-II for samples treated as in (A) as indicated above the blot. Lower blot shows total H3 loading control. For each blot, one of five biological replicates is shown. **(C)** Flow cytometric analysis of autophagosomes with CYTO-ID assay (Enzo) at day-3 +24h, +48h, and +72h in ABCs differentiating under standard conditions or in the presence of DMSO (vehicle) or UNC0638 (2 μM). Autophagy in Std, DMSO and UNC0638 treatment conditions is represented in different shades of grey, from darker to lighter, respectively. Data is representative of 3 biological replicates from a single experiment. **(D)** Flow cytometric analysis of surface phenotype of differentiating populations at day-6 generated under standard conditions (std) or in the presence of DMSO (vehicle) or UNC0638 (2 μM) plots showing CD19 vs CD20 (left) and CD38 vs CD138 (right). Percentages of total cell populations are shown following single cells and viability discrimination with gates established using matching isotype controls. One of six replicates from three independent experiments is shown. **(E)** Representative density flow plots showing gating strategy for cell division generations determined from CFSE dilution at 24, 48 and 72h of the ABC to plasmablast transition under standard conditions (left panel). Matching bar charts showing the frequency (% of cells/generation) at 24, 48 and 72h of the ABC to plasmablast transition for Std, DMSO and UNC0638 conditions. Bars represents mean ± SD of 3 replicates from a single experiment. *, P < 0.03 (Paired t-test) (right panel). **(F)** ELIspot results for IgM and IgG secretion for day-6 cells generated under standard, DMSO (vehicle control) or UNC0638 treated conditions. Representative wells shown on left and matching bar chart showing the spot counts (right). Bars are mean ± SD of 3 replicates from 2 independent experiments.

## References

1 Nutt, S. L., Hodgkin, P. D., Tarlinton, D. M. & Corcoran, L. M. The generation of antibody-secreting plasma cells. Nat Rev Immunol 15, 160–171, doi:10.1038/nri3795 (2015).

2 Scharer, C. D., Barwick, B. G., Guo, M., Bally, A. P. R. & Boss, J. M. Plasma cell differentiation is controlled by multiple cell division-coupled epigenetic programs. Nat Commun 9, 1698, doi:10.1038/s41467-018-04125-8 (2018).

3 Jourdan, M. et al. Characterization of a transitional preplasmablast population in the process of human B cell to plasma cell differentiation. Journal of Immunology 187, 3931–3941, doi:10.4049/jimmunol.1101230 (2011).

4 Gilchrist, M. et al. Systems biology approaches identify ATF3 as a negative regulator of Toll-like receptor 4. Nature 441, 173–178, doi:nature04768 [pii] 10.1038/nature04768 (2006).

5 Stephenson, S. et al. Growth Factor-like Gene Regulation Is Separable from Survival and Maturation in Antibody-Secreting Cells. J Immunol 202, 1287–1300, doi:10.4049/jimmunol.1801407 (2019).

6 Davidson, E. H. The regulatory genome : gene regulatory networks in development and evolution. (Academic, 2006).

7 Rothenberg, E. V. Causal Gene Regulatory Network Modeling and Genomics: Second-Generation Challenges. Journal of Computational Biology 26, 703–718, doi:10.1089/cmb.2019.0098 (2019).

8 Ramsey, S. A. et al. Epigenome-Guided Analysis of the Transcriptome of Plaque Macrophages during Atherosclerosis Regression Reveals Activation of the Wnt Signaling Pathway. Plos Genetics 10, doi:ARTN e1004828 10.1371/journal.pgen.1004828 (2014).

9 Sciammas, R. et al. Graded expression of interferon regulatory factor-4 coordinates isotype switching with plasma cell differentiation. Immunity 25, 225–236 (2006).

10 Klein, U. et al. Transcription factor IRF4 controls plasma cell differentiation and class-switch recombination. Nat Immunol 7, 773–782 (2006).

11 Kallies, A. et al. Initiation of plasma-cell differentiation is independent of the transcription factor Blimp-1. Immunity 26, 555–566, doi:S1074-7613(07)00250-6 [pii] 10.1016/j.immuni.2007.04.007 (2007).

12 Lin, K. I., Angelin-Duclos, C., Kuo, T. C. & Calame, K. Blimp-1-dependent repression of Pax-5 is required for differentiation of B cells to immunoglobulin M-secreting plasma cells. Mol Cell Biol 22, 4771–4780 (2002).

13 Tellier, J. et al. Blimp-1 controls plasma cell function through the regulation of immunoglobulin secretion and the unfolded protein response. Nat Immunol 17, 323–330, doi:10.1038/ni.3348 (2016).

14 Taubenheim, N. et al. High rate of antibody secretion is not integral to plasma cell differentiation as revealed by XBP-1 deficiency. J Immunol 189, 3328–3338, doi:10.4049/jimmunol.1201042 (2012).

15 Hu, C. C., Dougan, S. K., McGehee, A. M., Love, J. C. & Ploegh, H. L. XBP-1 regulates signal transduction, transcription factors and bone marrow colonization in B cells. Embo J 28, 1624–1636, doi:emboj2009117 [pii] 10.1038/emboj.2009.117 (2009).

16 Tirosh, B., Iwakoshi, N. N., Glimcher, L. H. & Ploegh, H. L. XBP-1 specifically promotes IgM synthesis and secretion, but is dispensable for degradation of glycoproteins in primary B cells. J Exp Med 202, 505–516 (2005).

17 Minnich, M. et al. Multifunctional role of the transcription factor Blimp-1 in coordinating plasma cell differentiation. Nat Immunol 17, 331–343, doi:10.1038/ni.3349 (2016).

18 Pasqualucci, L. et al. Inactivation of the PRDM1/BLIMP1 gene in diffuse large B cell lymphoma. J Exp Med 203, 311–317 (2006).

19 Mandelbaum, J. et al. BLIMP1 is a tumor suppressor gene frequently disrupted in activated B cell-like diffuse large B cell lymphoma. Cancer Cell 18, 568–579, doi:10.1016/j.ccr.2010.10.030 (2010).

20 Calado, D. P. et al. Constitutive canonical NF-kappaB activation cooperates with disruption of BLIMP1 in the pathogenesis of activated B cell-like diffuse large cell lymphoma. Cancer Cell 18, 580–589, doi:10.1016/j.ccr.2010.11.024 (2010).

21 Bikoff, E. K., Morgan, M. A. & Robertson, E. J. An expanding job description for Blimp-1/PRDM1. Curr Opin Genet Dev, doi:S0959-437X(09)00093-8 [pii] 10.1016/j.gde.2009.05.005 (2009).

22 Su, S. T. et al. Involvement of histone demethylase LSD1 in Blimp-1-mediated gene repression during plasma cell differentiation. Mol Cell Biol 29, 1421–1431, doi:MCB.01158-08 [pii] 10.1128/MCB.01158-08 (2009).

23 Yu, J., Angelin-Duclos, C., Greenwood, J., Liao, J. & Calame, K. Transcriptional repression by blimp-1 (PRDI-BF1) involves recruitment of histone deacetylase. Mol Cell Biol 20, 2592–2603 (2000).

24 Gyory, I., Wu, J., Fejer, G., Seto, E. & Wright, K. L. PRDI-BF1 recruits the histone H3 methyltransferase G9a in transcriptional silencing. Nat Immunol 5, 299–308 (2004).

25 Guo, M. et al. EZH2 Represses the B Cell Transcriptional Program and Regulates Antibody-Secreting Cell Metabolism and Antibody Production. J Immunol 200, 1039–1052, doi:10.4049/jimmunol.1701470 (2018).

26 Mochizuki, K. et al. Repression of Somatic Genes by Selective Recruitment of HDAC3 by BLIMP1 Is Essential for Mouse Primordial Germ Cell Fate Determination. Cell Rep 24, 2682–2693 e2686, doi:10.1016/j.celrep.2018.07.108 (2018).

27 Ancelin, K. et al. Blimp1 associates with Prmt5 and directs histone arginine methylation in mouse germ cells. Nat Cell Biol 8, 623–630 (2006).

28 Mozzetta, C., Boyarchuk, E., Pontis, J. & Ait-Si-Ali, S. Sound of silence: the properties and functions of repressive Lys methyltransferases. Nat Rev Mol Cell Biol 16, 499–513, doi:10.1038/nrm4029 (2015).

29 Kouzarides, T. Chromatin modifications and their function. Cell 128, 693–705, doi:10.1016/j.cell.2007.02.005 (2007).

30 Arpin, C. et al. Generation of memory B cells and plasma cells in vitro. Science 268, 720–722, doi:10.1126/science.7537388 (1995).

31 Cocco, M. et al. In vitro generation of long-lived human plasma cells. J Immunol 189, 5773–5785, doi:10.4049/jimmunol.1103720 (2012).

32 Care, M. A., Westhead, D. R. & Tooze, R. M. Parsimonious Gene Correlation Network Analysis (PGCNA): a tool to define modular gene co-expression for refined molecular stratification in cancer. NPJ Syst Biol Appl 5, 13, doi:10.1038/s41540-019-0090-7 (2019).

33 Monti, S. et al. Molecular profiling of diffuse large B-cell lymphoma identifies robust subtypes including one characterized by host inflammatory response. Blood 105, 1851–1861 (2005).

34 Shaffer, A. L. et al. A library of gene expression signatures to illuminate normal and pathological lymphoid biology. Immunol Rev 210, 67–85, doi:IMR373 [pii] 10.1111/j.0105-2896.2006.00373.x (2006).

35 Subramanian, A. et al. Gene set enrichment analysis: a knowledge-based approach for interpreting genome-wide expression profiles. Proc Natl Acad Sci U S A 102, 15545–15550 (2005).

36 Culhane, A. C. et al. GeneSigDB--a curated database of gene expression signatures. Nucleic acids research 38, D716–725, doi:10.1093/nar/gkp1015 (2010).

37 Rosenwald, A. et al. Molecular diagnosis of primary mediastinal B cell lymphoma identifies a clinically favorable subgroup of diffuse large B cell lymphoma related to Hodgkin lymphoma. The Journal of experimental medicine 198, 851–862, doi:10.1084/jem.20031074 (2003).

38 Doody, G. M., Stephenson, S. & Tooze, R. M. BLIMP-1 is a target of cellular stress and downstream of the unfolded protein response. Eur J Immunol 36, 1572–1582 (2006).

39 Doody, G. M., Stephenson, S., McManamy, C. & Tooze, R. M. PRDM1/BLIMP-1 modulates IFN-gamma-dependent control of the MHC class I antigen-processing and peptide-loading pathway. Journal of immunology 179, 7614–7623 (2007).

40 Doody, G. M. et al. An extended set of PRDM1/BLIMP1 target genes links binding motif type to dynamic repression. Nucleic Acids Res 38, 5336–5350, doi:gkq268 [pii] 10.1093/nar/gkq268 (2010).

41 Langmead, B. & Salzberg, S. L. Fast gapped-read alignment with Bowtie 2. Nature methods 9, 357–359, doi:10.1038/nmeth.1923 (2012).

42 Guo, Y., Mahony, S. & Gifford, D. K. High resolution genome wide binding event finding and motif discovery reveals transcription factor spatial binding constraints. PLoS computational biology 8, e1002638, doi:10.1371/journal.pcbi.1002638 (2012).

43 Zhang, Y. et al. Model-based Analysis of ChIP-Seq (MACS). Genome biology 9, doi:ARTN R137 10.1186/gb-2008-9-9-r137 (2008).

44 Heinz, S. et al. Simple combinations of lineage-determining transcription factors prime cis-regulatory elements required for macrophage and B cell identities. Molecular cell 38, 576–589, doi:10.1016/j.molcel.2010.05.004 (2010).

45 Finkin, S., Hartweger, H., Oliveira, T. Y., Kara, E. E. & Nussenzweig, M. C. Protein Amounts of the MYC Transcription Factor Determine Germinal Center B Cell Division Capacity. Immunity 51, 324-+, doi:10.1016/j.immuni.2019.06.013 (2019).

46 Luo, W., Weisel, F. & Shlomchik, M. J. B Cell Receptor and CD40 Signaling Are Rewired for Synergistic Induction of the c-Myc Transcription Factor in Germinal Center B Cells. Immunity 48, 313-+, doi:10.1016/j.immuni.2018.01.008 (2018).

47 Kuo, T. C. & Calame, K. L. B lymphocyte-induced maturation protein (Blimp)-1, IFN regulatory factor (IRF)-1, and IRF-2 can bind to the same regulatory sites. J Immunol 173, 5556–5563 (2004).

48 Doody, G. M., Stephenson, S., McManamy, C. & Tooze, R. M. PRDM1/BLIMP-1 modulates IFN-gamma-dependent control of the MHC class I antigen-processing and peptide-loading pathway. J Immunol 179, 7614–7623 (2007).

49 Yamamoto, K., Yoshida, H., Kokame, K., Kaufman, R. J. & Mori, K. Differential contributions of ATF6 and XBP1 to the activation of endoplasmic reticulum stress-responsive cis-acting elements ERSE, UPRE and ERSE-II. Journal of Biochemistry 136, 343–350, doi:10.1093/jb/mvh122 (2004).

50 Kundaje, A. et al. Ubiquitous heterogeneity and asymmetry of the chromatin environment at regulatory elements. Genome Research 22, 1735–1747, doi:10.1101/gr.136366.111 (2012).

51 Pengo, N. et al. Plasma cells require autophagy for sustainable immunoglobulin production. Nature immunology 14, 298–305, doi:10.1038/ni.2524 (2013).

52 Martinez-Martin, N. et al. A switch from canonical to noncanonical autophagy shapes B cell responses. Science 355, 641–647, doi:10.1126/science.aal3908 (2017).

53 Shin, H. M. et al. Epigenetic Modifications Induced by Blimp-1 Regulate CD8(+) T Cell Memory Progression during Acute Virus Infection. Immunity 39, 661–675, doi:10.1016/j.immuni.2013.08.032 (2013).

54 Care, M. A. et al. A Microarray Platform-Independent Classification Tool for Cell of Origin Class Allows Comparative Analysis of Gene Expression in Diffuse Large B-cell Lymphoma. PLoS ONE 8, e55895, doi:10.1371/journal.pone.0055895 (2013).

55 Carotta, S. et al. The transcription factors IRF8 and PU.1 negatively regulate plasma cell differentiation. Journal of Experimental Medicine 211, 2169–2181, doi:10.1084/jem.20140425 (2014).

56 Willis, S. N. et al. Environmental sensing by mature B cells is controlled by the transcription factors PU.1 and SpiB. Nat Commun 8, doi:ARTN 1426 10.1038/s41467-017-01605-1 (2017).

57 Wang, H. et al. Transcription factors IRF8 and PU.1 are required for follicular B cell development and BCL6-driven germinal center responses. P Natl Acad Sci USA 116, 9511–9520, doi:10.1073/pnas.1901258116 (2019).

58 Lenz, G. et al. Molecular subtypes of diffuse large B-cell lymphoma arise by distinct genetic pathways. P Natl Acad Sci USA 105, 13520–13525, doi:10.1073/pnas.0804295105 (2008).

59 Care, M. A. et al. SPIB and BATF provide alternate determinants of IRF4 occupancy in diffuse large B-cell lymphoma linked to disease heterogeneity. Nucleic Acids Research 42, 7588–7610, doi:10.1093/nar/gku451 (2014).

60 Iwakoshi, N. N. et al. Plasma cell differentiation and the unfolded protein response intersect at the transcription factor XBP-1. Nat Immunol 4, 321–329 (2003).

61 Iwakoshi, N. N., Lee, A. H. & Glimcher, L. H. The X-box binding protein-1 transcription factor is required for plasma cell differentiation and the unfolded protein response. Immunol Rev 194, 29–38 (2003).

62 Todd, D. J. et al. XBP1 governs late events in plasma cell differentiation and is not required for antigen-specific memory B cell development. J Exp Med, doi:jem.20090738 [pii] 10.1084/jem.20090738 (2009).

63 Al-Maskari, M. et al. Site-1 protease function is essential for the generation of antibody secreting cells and reprogramming for secretory activity. Sci Rep 8, doi:ARTN 14338 10.1038/s41598-018-32705-7 (2018).

64 Gass, J. N., Jiang, H. Y., Wek, R. C. & Brewer, J. W. The unfolded protein response of B-lymphocytes: PERK-independent development of antibody-secreting cells. Mol Immunol 45, 1035–1043, doi:S0161-5890(07)00655-4 [pii] 10.1016/j.molimm.2007.07.029 (2008).

65 Cheung, K. L. et al. Distinct Roles of Brd2 and Brd4 in Potentiating the Transcriptional Program for Th17 Cell Differentiation. Molecular Cell 65, 1068-+, doi:10.1016/j.molcel.2016.12.022 (2017).

66 Scheer, S. & Zaph, C. The Lysine Methyltransferase G9a in Immune Cell Differentiation and Function. Front Immunol 8, doi:ARTN 429 10.3389/fimmu.2017.00429 (2017).

67 Brown, P. J. & Muller, S. Open access chemical probes for epigenetic targets. Future Med Chem 7, 1901–1917, doi:10.4155/fmc.15.127 (2015).

68 Vedadi, M. et al. A chemical probe selectively inhibits G9a and GLP methyltransferase activity in cells. Nat Chem Biol 7, 566–574, doi:10.1038/nchembio.599 (2011).

69 Sato, T. et al. Transcriptional Selectivity of Epigenetic Therapy in Cancer. Cancer Res 77, 470–481, doi:10.1158/0008-5472.CAN-16-0834 (2017).

70 Shankar, S. R. et al. G9a, a multipotent regulator of gene expression. Epigenetics 8, 16–22, doi:10.4161/epi.23331 (2013).

