## Supplemental Methods for "A dichotomy in association of core transcription factors and gene regulation during the activated B-cell to plasmablast transition"

### Contents

Gene expression analysis
  Expression data-sets
  Normalisation and analysis
PGCNA networks
  Background
  Data preparation
  PGCNA2 network generation
ChIP-seq analysis
  Data-sets
  Alignment and peak discovery
  Peak overlap and filtering
  Peak annotation
  Motif discovery
Data visualisation
  Network visualisation
  Heatmaps
  ChIP-seq clustering and visualisation
  Graphs
  Venn diagrams
  Motif logos
Statistical analyses
  Gene signature data
  Enrichment analysis
Data processing
Data and software availability
References

### Gene expression analysis

#### Expression data-sets

Two gene expression data-sets were generated, a total B cell (totalB) and a negatively selected (CD23 depletion) memory B cell set (memB; referred to as m. in figures), both from 3 healthy donors, from day-0 (resting B-cell; totalB only) on day-3 after activation with CD40L/anti-BCR/cytokines (ABC) and then at intervals of 3h, 6h, 12h, 24h, 48h and 72h (PB) after transition into conditions (continued IL2 and IL21 only) that support the ABC/PB transition.  A G9A inhibited data-set (G9A_i) was generated for the totalB data with the inclusion of the small molecule inhibitor UNC0638 at day-3, with samples taken at the same intervals as the standard conditions.

#### Normalisation and analysis

Illumina GenomeStudio Gene Expression Module was employed for initial data processing followed by the R Lumi package (v2.36.0).[1] Probes not detected on three or more arrays were removed and the remaining data variance-stabilizing transformed (VST) and quantile normalized. A linear model was fitted to the gene expression data using the R Limma package (v3.40.6).[2] Differentially expressed genes between the Standard/G9A_i contrasts were gauged using the Limma empirical Bayes statistics module, adjusting for multiple testing using Benjamini and Hochberg correction. Probes were re-annotated using the MyGene.info (http://mygene.info) API using all available references (e.g. NCBI Entrez, Ensembl etc.) and any ambiguous mappings manually assigned.[3] Finally, probes for each gene were merged by taking the median value for probe sets with a Pearson correlation ≥0.2 and the maximum value for those with a correlation <0.2 (as used by Monti *et al*).[4]

### PGCNA networks

#### Background

For details and validation of the Parsimonious Gene Correlation Network Analysis (PGCNA) approach see our other work.[5] Here a brief description of the method will be given. After informative genes are selected they are used to calculate Spearman's rank correlations for all gene pairs using the Python scipy.stats package. For each gene (row) in a correlation matrix only the 3 most correlated edges per gene are retained. The resulting matrix $M$, with entries written as $M = (m_{ij})$ is made symmetrical by setting $m_{ij} = m_{ji}$ for all indices $i$ and $j$ so that $M = \mathrm{M}^{\mathrm{T}}$ (its transpose). The correlation matrices are clustered using a community detection algorithm (Louvain v0.3 or Leidenalg v0.7.0; see PGCNA2) and the 100 best (judged by modularity score) used for downstream analysis.[6,7]

#### Data preparation

The totalB and memB standard condition time course data were used for PGCNA network construction. Before probes were merged per-gene the most informative probes were selected for the different time courses. All probes with a $\sigma^2 > 0.025$ (across median values per time point) were selected, giving 11,418 and 10,183 probes for the totalB and memB sets respectively. The selected probes were used to filter the complete (not merged across donors) data-sets and finally the probes merged per gene. This gave 9,362 and 8,431 genes for the totalB and memB data-sets.

#### PGCNA2 network generation

For this project the community detection method used by PGCNA (Louvain/FastUnfold) was replaced with an improved algorithm that guarantees well-connected communities (Leidenalg).[6,7] In order to test if the Leidenalg improved the results of the PGCNA approach it was compared to the Louvain method for the BRCA/COAD data-sets (used in the original PGCNA paper) and the totalB and memB data-sets (used here) using a Spearman correlation matrix reduced to 3 edges per gene (EPG).[8] For the BRCA/COAD analysis the

Louvain results were compared to 3 rounds of Leidenalg runs, whilst for the smaller totalB/memB data-sets the Louvain results were compared to 6 rounds of Leidenalg runs. For the Louvain method each round consisted of 10,000 iterations, whilst for Leidenalg each round consisted of 1,000 iterations (with optimiser set to iterate until no partition improvement). For both Louvain/Leidenalg for each round the top 100 results (judged by modularity score) were then analysed for biological enrichment against a signature database (see Gene signature data) by generating Scaled cluster enrichment scores (see original PGCNA methods for details).[5] Methods Figure 1 and 2 show that in 3 of the 4 tested data-sets Leidenalg improved the biological enrichment. The optimal Leidenalg results were then used for downstream visualisation and analysis. This resulted in two networks: PGCNA-totalB (9,362 genes, 22,921 edges) and PGCNA-memB (8,431 genes, 20,524 edges).

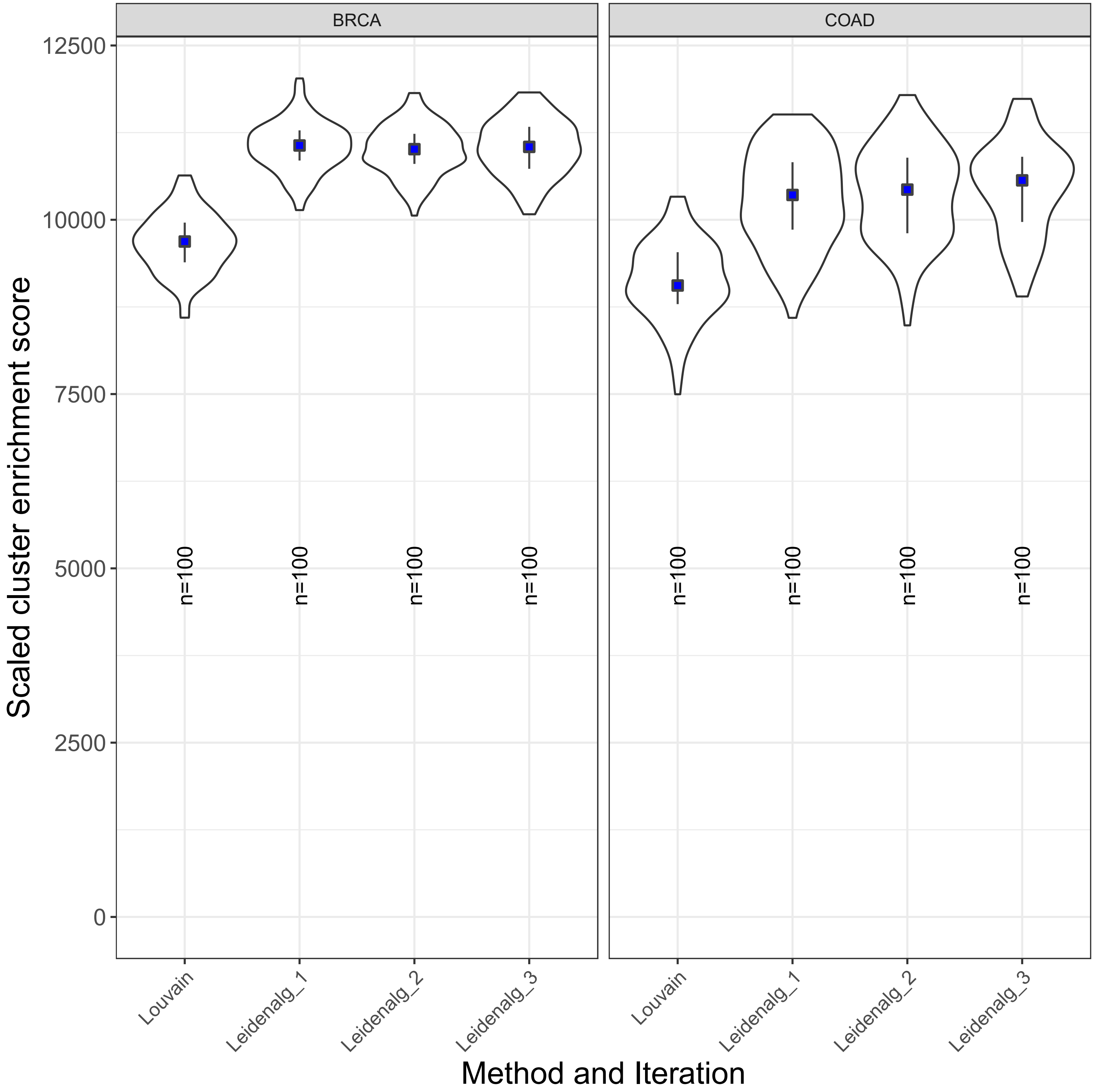


**Methods Fig 1. BRCA/COAD.** Enrichment of gene ontology and signatures was assessed using a scaled cluster enrichment score (SCES) and compared between data clustered with Louvain (FastUnfold) or Leidenalg for parsimonious matrices with edges per gene (EPG) of 3. Violin plots display the distribution along with median (blue square) and the IQR.

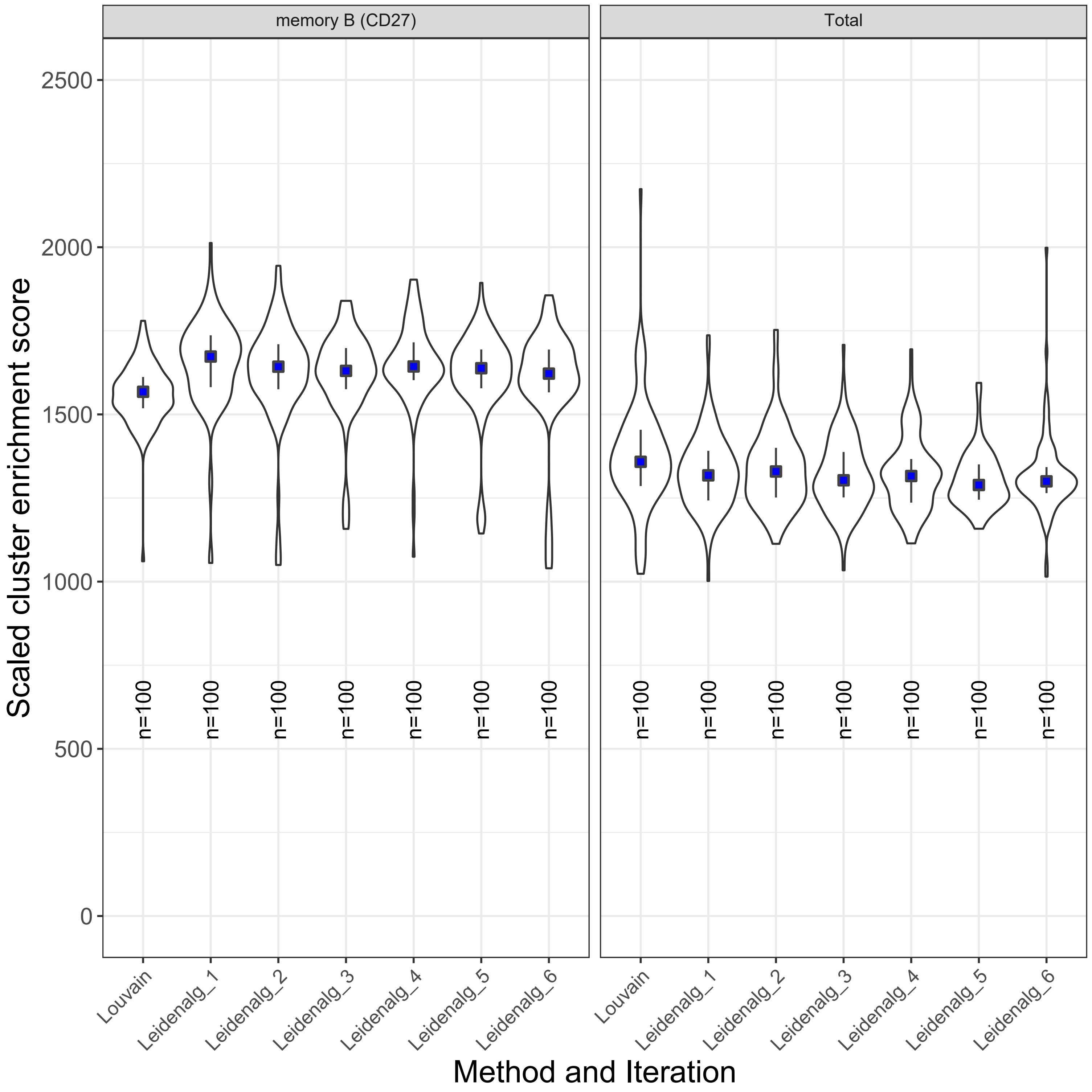


**Methods Fig 2. memory/Total.** Enrichment of gene ontology and signatures was assessed using a scaled cluster enrichment score (SCES) and compared between data clustered with Louvain (FastUnfold) or Leidenalg for parsimonious matrices with edges per gene (EPG) of 3. Violin plots display the distribution along with median (blue square) and the IQR.

### ChIP-seq analysis

#### Data-sets

| File | TF | Condition | TotalReads | TotalAligned | %Aligned |
|---|---|---|---|---|---|
| Day6PB-B_XBP1_Std | XBP1 | Std | 26,511,959 | 21,722,657 | 81.94 |
| Day6PB-B_Input_Std | - | Std | 29,074,912 | 23,855,970 | 82.05 |
| Day6PB-memB_BLIMP1_Std | BLIMP1 | Std | 35,002,987 | 29,416,625 | 84.04 |
| Day6PB-memB_CTCF_Std | CTCF | Std | 36,105,839 | 31,556,061 | 87.40 |
| Day6PB-memB_H3K27Ac_Std | H3K27ac | Std | 54,433,687 | 47,458,206 | 87.19 |
| Day6PB-memB_H3K27me3_Std | H3K27me3 | Std | 52,430,237 | 45,213,387 | 86.24 |
| Day6PB-memB_H3K4me3_Std | H3K4me3 | Std | 53,132,933 | 43,282,575 | 81.46 |
| Day6PB-memB_H3K9me2_Std | H3K9me2 | Std | 43,552,007 | 36,498,541 | 83.80 |
| Day6PB-memB_IRF4_Std | IRF4 | Std | 34,346,524 | 28,691,545 | 83.54 |
| Day6PB-memB_Input_Std | - | Std | 26,944,620 | 14,681,441 | 54.49 |
| Day6PB-memB_BLIMP1_0638 | BLIMP1 | UNC0638 | 26,740,354 | 22,690,984 | 84.86 |
| Day6PB-memB_H3K27Ac_0638 | H3K27ac | UNC0638 | 39,826,235 | 34,929,002 | 87.70 |
| Day6PB-memB_H3K27me3_0638 | H3K27me3 | UNC0638 | 36,296,186 | 30,987,431 | 85.37 |
| Day6PB-memB_H3K4me3_0638 | H3K4me3 | UNC0638 | 40,544,784 | 33,485,297 | 82.59 |
| Day6PB-memB_H3K9me2_0638 | H3K9me2 | UNC0638 | 42,618,182 | 34,930,038 | 81.96 |
| Day6PB-memB_Input_0638 | - | UNC0638 | 21,062,154 | 11,530,118 | 54.74 |

#### Alignment and peak discovery

Before aligning, reads were trimmed to remove adapters and low confidence regions using a python script. A 4 base sliding window was run along each read, if the average Q Phred score for a window was < 20 the read was trimmed at the window start, finally any match to adapter sequences were trimmed.

Trimmed reads were aligned to GRCh38 (UCSC analysis set) using Bowtie2 (v2.3.5.1) with the --very-sensitive parameter.[9] The resultant SAM files were converted to BAM using samtools (v1.9) with the quality filter set to 20 (-q 20).[10] The BAM files were analysed for peaks using GEM (v3.4), with quality filter set to 1 (-q 1) and MACS2 (v2.1.2) with quality filter set to 0.05 (-q 0.05).[11,12]

The resultant BAM files were converted to BED files and read cross-correlation was assessed using MaSC (v1.2.1).[13] Reads were extended to the estimated fragment length, and a scaled (reads per million; rpm) BED file generated. This was converted to a coverage file using the UCSC genomeCoverageBed tool and then to a BigWig file using UCSC bedGraphToBigWig for peak visualisation.[14]

#### Peak overlap and filtering

ChIP-seq peak overlaps were used in two contexts. Firstly, to reduce false positive peak discovery a high-confidence merged peak set was created by overlapping the GEM/MACS2 peaks. Secondly, the resultant high-confidence peaks were used to find the overlap between the different transcription factors (TFs) creating a non-redundant set of binding regions. The peak centres for all data-sets were ordered per chromosome.

Starting at the beginning of each chromosome peaks were added to a cluster. New peaks were only added to the cluster if the distance between them and the first peak was ≤ 500 bp, else a new cluster was started. As new clusters were generated if peaks in the earlier cluster were closer to the new cluster's centre they were moved into the new cluster (thus a peak can only belong to one cluster). The minimum GEM/MACS2 $-log_{10}Q$ for a peak to start a new cluster was set to ≥ 5, however, this was lowered to 1 for the addition of peaks to existing clusters. Note that the number of peaks in the TF overlap sets are less than the individual peaks per TF due to the merging of some close peaks. This gave the final individual high-confidence sets: BLIMP1 (n=4,244), IRF4 (n=9,259) and XBP1 (n=602) along with the Union set consisting of all peaks in the overlap of BLIMP1_IRF4_XBP1 (n=11,467). The results from peak merging and overlap analysis can be found in Supplemental Table 5.

#### Peak annotation

Version 28 of the Gencode gene annotation data was downloaded from UCSC, the genes were re-annotated using the HUGO Gene Nomenclature Committee annotations (2018/06/08 version).[14,15] Each peak was assigned the nearest gene (TSS) as its primary gene. In addition annotatePeaks.pl from the HOMER suite (v4.11.1) was used to provide detailed peak annotations (Promoter: -1kb – 100bp, TTS: -100bp – 1kb, Exonic/Intronic: > 100bp from Promoter/TTS within gene, Intergenic: >1kb from Promoter/TTS outside gene).[16]

#### Motif discovery

BED files were generated for the peak overlap sets, -/+ 125 bases around each peak centre (or overlap cluster centre). These were analysed for *de novo* motifs of length 8 – 14 using findMotifsGenome.pl from the HOMER suite.

### Data visualisation

#### Network visualisation

The optimal PGCNA-totalB/PGCNA-memB networks were converted to a list of edges and nodes and uploaded into the Gephi package (version 0.9.2).[17] Degree and Betweenness Centrality were calculated, and the latter used to adjust node sizes. The network layout was generated using the ForceAtlas2 approach, and interactive HTML5 web visualizations exported using the sigma.js library (https://github.com/oxfordinternetinstitute/gephi-plugins/tree/sigmaexporter-plugin). The interactive visualisations can be found at https://mcare.link/g9a.

#### Heatmaps

The gene expression data and GSE results were both visualised using the Broad GENE-E package (https://software.broadinstitute.org/GENE-E/). For visualisation of expression data, the timepoint median expression values were row-normalised (z-scores), while for GSE visualisation the signature enrichment/depletion z-scores were used. In both cases the data was hierarchically clustered (Pearson correlations and average linkage).

#### ChIP-seq clustering and visualisation

The high-confidence peak sets for BLIMP1, IRF4 and XBP1 along with the Union set (overlap of individual high-confidence BLIMP1, IRF4 and XBP1 peaks; see supplemental Table 5) were analysed using the deepTools2 suite (v3.3.0).[18] Using bamCoverage peaks were normalised to bins per million mapped reads (BPM) and extended to their MaSC estimated fragment length (e.g. --normalizeUsing BPM --extendReads 140 --binSize 10). Scores per region were calculated with computeMatrix using a BED file reference for a ± 1000bp region (--referencePoint center -b 1000 -a 1000 --skipZeros). The resulting matrix was k-means clustered and then visualised using plotHeatmap (--kmeans 6).

#### Graphs

All graph visualisations were generated using the R package ggplot2 (v3.2.1), with the viridis (v0.5.1) colour-blind friendly colour scheme.[19]

#### Venn diagrams

Venn diagrams were generated using the R package VennDiagram (v1.6.20).

#### Motif logos

The *de novo* motifs generated by HOMER were converted to information scaled pdfs using the python API for the WebLogo package (v3.6.0).[20]

### Statistical analyses

#### Gene signature data

A data-set of 17,904 gene signatures was created by merging signatures downloaded from http://lymphochip.nih.gov/signaturedb/ (SignatureDB), http://www.broadinstitute.org/gsea/msigdb/index.jsp MSigDB V6.2 (MSigDB C1–C7 and H; excluding C5. With MIPS signatures from version 3.1 and PID signatures from version 4 added back), http://compbio.dfci.harvard.edu/genesigdb/ Gene Signature Database V4 (GeneSigDB), UniProt keywords (parsed XML from http://www.uniprot.org/downloads), and fifteen papers.[4,21,30–37,22–29] A gene ontology gene set was created using an in-house python script. This parses a gene association file (http://geneontology.org/page/download-go-annotations) to link genes with ontology terms and then uses the ontology structure (.obo file; http://purl.obolibrary.org/obo/go.obo) to propagate these terms up to the root. The resultant gene set contained 22,782 terms. The gene-ontology and gene-signatures sets were merged to give a final signature set of 40,686 terms.

#### Enrichment analysis

The gene signature enrichment (GSE) was assessed using a hypergeometric test, in which the draw is the gene list genes, the successes are the signature genes, and the population is the genes present on the platform. The resultant p-values are then adjusted for multiple testing using Benjamini and Hochberg correction. GSE was carried out in three different contexts: to assess the biological enrichment of the PGCNA-totalB/PGCNA-memB network modules, to compare the TF bound genes against the PGCNA-memB network and finally to compare the G9A_i differentially expressed genes against the PGCNA-memB network.

For the PGCNA network GSE analyses the genes per module were compared against the 40,686-signature database (background: all the genes in that network). For analysis of TF bound genes against the PGCNA-memB modules the peaks were assigned to the nearest TSS (≤10kb), the resultant genes were then compared against the genes per PGCNA-memB module (background: PGCNA-memB genes). Finally, for the G9A_i enrichment assessment the 72 hours differentially expressed genes (FDR < 0.05, FC > 1.2) were compared against the genes per PGCNA-memB module (background: PGCNA-memB genes).

#### Data processing

All analyses were undertaken on MARC1, part of the High Performance Computing and Leeds Institute for Data Analytics (LIDA) facilities at the University of Leeds, UK

#### Data and software availability

Interactive networks and all meta-data are available at https://mcare.link/g9a.  PGCNA python scripts are available at https://github.com/medmaca/PGCNA.

#### References


1. Du, P., Kibbe, W. a & Lin, S. M. lumi: a pipeline for processing Illumina microarray. *Bioinformatics* 24, 1547–8 (2008).

2. Ritchie, M. E. *et al.* limma powers differential expression analyses for RNA-sequencing and microarray studies. *Nucleic Acids Res.* 43, e47 (2015).

3. Xin, J. *et al.* High-performance web services for querying gene and variant annotation. *Genome Biol.* 17, 91 (2016).

4. Monti, S. *et al.* Molecular profiling of diffuse large B-cell lymphoma identifies robust subtypes including one characterized by host inflammatory response. *Blood* 105, 1851–61 (2005).

5. Care, M. A., Westhead, D. R. & Tooze, R. M. Parsimonious Gene Correlation Network Analysis (PGCNA): a tool to define modular gene co-expression for refined molecular stratification in cancer. *NPJ Syst. Biol. Appl.* 5, 13 (2019).

6. Traag, V. A., Waltman, L. & van Eck, N. J. From Louvain to Leiden: guaranteeing well-connected communities. *Sci. Rep.* 9, (2019).

7. Blondel, V. D., Guillaume, J.-L., Lambiotte, R. & Lefebvre, E. Fast unfolding of communities in large networks. *J. Stat. Mech. Theory Exp.* 2008, P10008 (2008).

8. Care, M. A., Westhead, D. R. & Tooze, R. M. Parsimonious Gene Correlation Network Analysis (PGCNA): a tool to define modular gene co-expression for refined molecular stratification in cancer. *NPJ Syst. Biol. Appl.* 5, 13 (2019).

9. Langmead, B. & Salzberg, S. L. Fast gapped-read alignment with Bowtie 2. *Nat. Methods* 9, 357–9 (2012).

10. Li, H. *et al.* The Sequence Alignment/Map format and SAMtools. *Bioinformatics* 25, 2078–9 (2009).

11. Guo, Y., Mahony, S. & Gifford, D. K. High resolution genome wide binding event finding and motif discovery reveals transcription factor spatial binding constraints. *PLoS Comput. Biol.* 8, e1002638 (2012).

12. Zhang, Y. *et al.* Model-based analysis of ChIP-Seq (MACS). *Genome Biol.* 9, R137 (2008).

13. Ramachandran, P., Palidwor, G. a, Porter, C. J. & Perkins, T. J. MaSC: mappability-

sensitive cross-correlation for estimating mean fragment length of single-end short-read sequencing data. *Bioinformatics* 29, 444–50 (2013).

14. Rhead, B. *et al.* The UCSC Genome Browser database: update 2010. *Nucleic Acids Res.* 38, D613-9 (2010).

15. Gray, K. a *et al.* Genenames.org: the HGNC resources in 2013. *Nucleic Acids Res.* 1–8 (2012). doi:10.1093/nar/gks1066

16. Heinz, S. *et al.* Simple Combinations of Lineage-Determining Transcription Factors Prime cis-Regulatory Elements Required for Macrophage and B Cell Identities. *Mol. Cell* 38, 576–589 (2010).

17. Bastian, M., Heymann, S. & Jacomy, M. Gephi : An Open Source Software for Exploring and Manipulating Networks. *ICWSM* 8, 361–362 (2009).

18. Ramírez, F. *et al.* deepTools2: a next generation web server for deep-sequencing data analysis. *Nucleic Acids Res.* 44, W160-5 (2016).

19. Wickham, H. ggplot2. *Wiley Interdiscip. Rev. Comput. Stat.* 3, 180–185 (2011).

20. Crooks, G. E., Hon, G., Chandonia, J.-M. & Brenner, S. E. WebLogo: a sequence logo generator. *Genome Res.* 14, 1188–90 (2004).

21. Subramanian, A. *et al.* Gene set enrichment analysis: a knowledge-based approach for interpreting genome-wide expression profiles. *Proc. Natl. Acad. Sci. U. S. A.* 102, 15545–50 (2005).

22. Culhane, A. C. *et al.* GeneSigDB--a curated database of gene expression signatures. *Nucleic Acids Res.* 38, D716-25 (2010).

23. Shaffer, A. L. *et al.* A library of gene expression signatures to illuminate normal and pathological lymphoid biology. *Immunol. Rev.* 210, 67–85 (2006).

24. Rosenwald, A. *et al.* Molecular diagnosis of primary mediastinal B cell lymphoma identifies a clinically favorable subgroup of diffuse large B cell lymphoma related to Hodgkin lymphoma. *J. Exp. Med.* 198, 851–862 (2003).

25. Compagno, M. *et al.* Mutations of multiple genes cause deregulation of NF-kappaB in diffuse large B-cell lymphoma. (Supplemental). *Nature* 459, 717–21 (2009).

26. Berry, M. P. R. *et al.* An interferon-inducible neutrophil-driven blood transcriptional signature in human tuberculosis. *Nature* 466, 973–977 (2010).

27. Ngo, V. N. *et al.* Oncogenically active MYD88 mutations in human lymphoma. *Nature* 470, 115–9 (2011).

28. Kidani, Y. *et al.* Sterol regulatory element-binding proteins are essential for the metabolic programming of effector T cells and adaptive immunity. *Nat. Immunol.* 14, 489–99 (2013).

29. Sadanandam, A. *et al.* A colorectal cancer classification system that associates cellular phenotype and responses to therapy. *Nat. Med.* 19, 619–25 (2013).

30. De Sousa E Melo, F. *et al.* Poor-prognosis colon cancer is defined by a molecularly distinct subtype and develops from serrated precursor lesions. *Nat. Med.* 19, 614–8 (2013).

31. Marisa, L. *et al.* Gene expression classification of colon cancer into molecular subtypes: characterization, validation, and prognostic value. *PLoS Med.* 10,

e1001453 (2013).

32. Masiero, M. *et al.* A core human primary tumor angiogenesis signature identifies the endothelial orphan receptor ELTD1 as a key regulator of angiogenesis. *Cancer Cell* 24, 229–41 (2013).

33. Bindea, G. *et al.* Spatiotemporal dynamics of intratumoral immune cells reveal the immune landscape in human cancer. *Immunity* 39, 782–795 (2013).

34. Huang, X. *et al.* Activation of the STAT3 signaling pathway is associated with poor survival in diffuse large B-cell lymphoma treated with R-CHOP. *J. Clin. Oncol.* 31, 4520–8 (2013).

35. Li, S. *et al.* Molecular signatures of antibody responses derived from a systems biology study of five human vaccines. *Nat. Immunol.* 15, 195–204 (2014).

36. Chiche, L. *et al.* Modular Transcriptional Repertoire Analyses of Adults With Systemic Lupus Erythematosus Reveal Distinct Type I and Type II Interferon Signatures. *Arthritis Rheumatol. (Hoboken, N.J.)* 66, 1583–95 (2014).

37. Newman, A. M. *et al.* Robust enumeration of cell subsets from tissue expression profiles. *Nat. Methods* 12, 1–10 (2015).
