## Supplementary figures and images for "A dichotomy in association of core transcription factors and gene regulation during the activated B-cell to plasmablast transition"

### Supplemental Figure 1

https://mcare.link/G9A_Paper_Total-Std-Mod

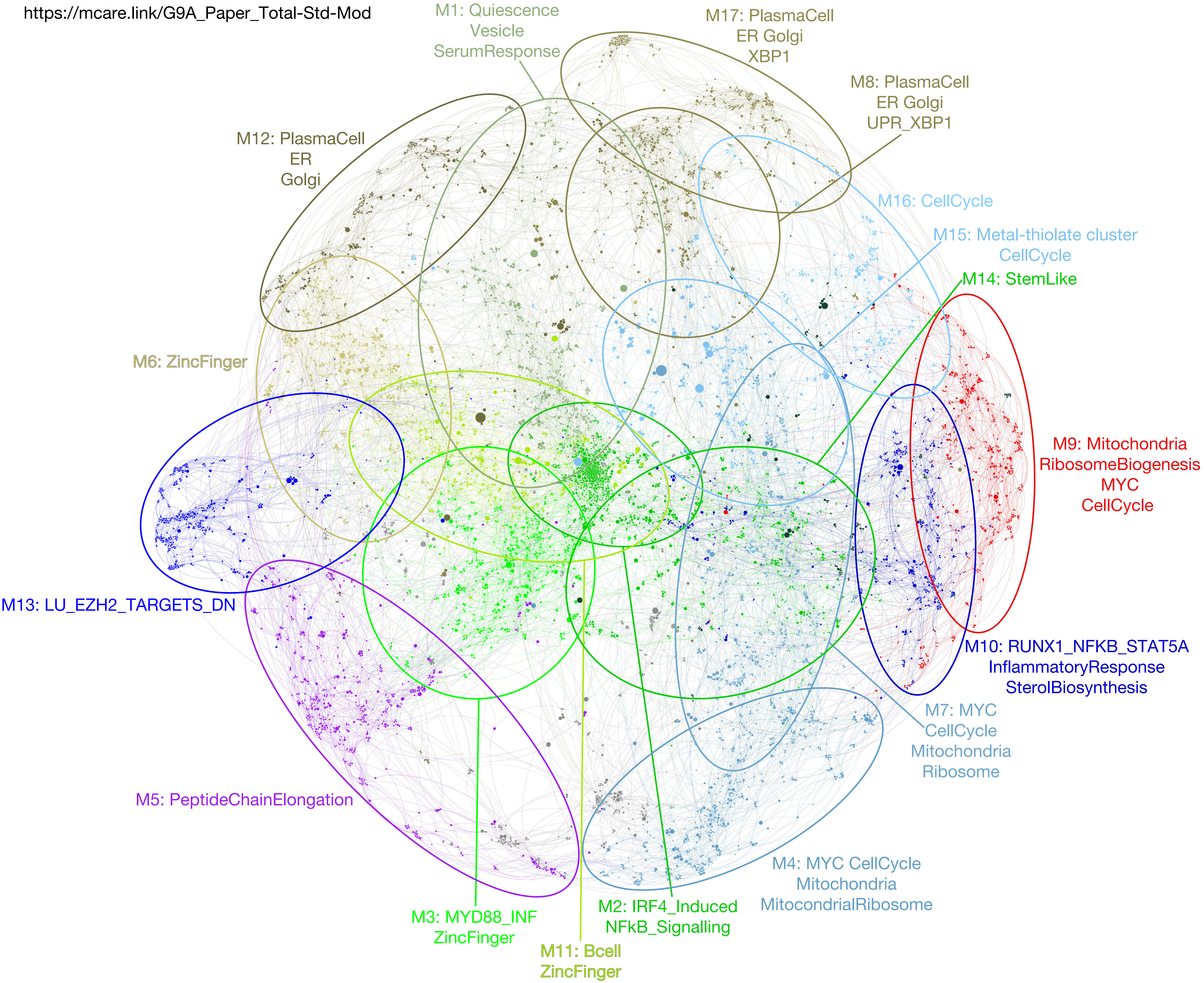

### Supplemental Figure 2

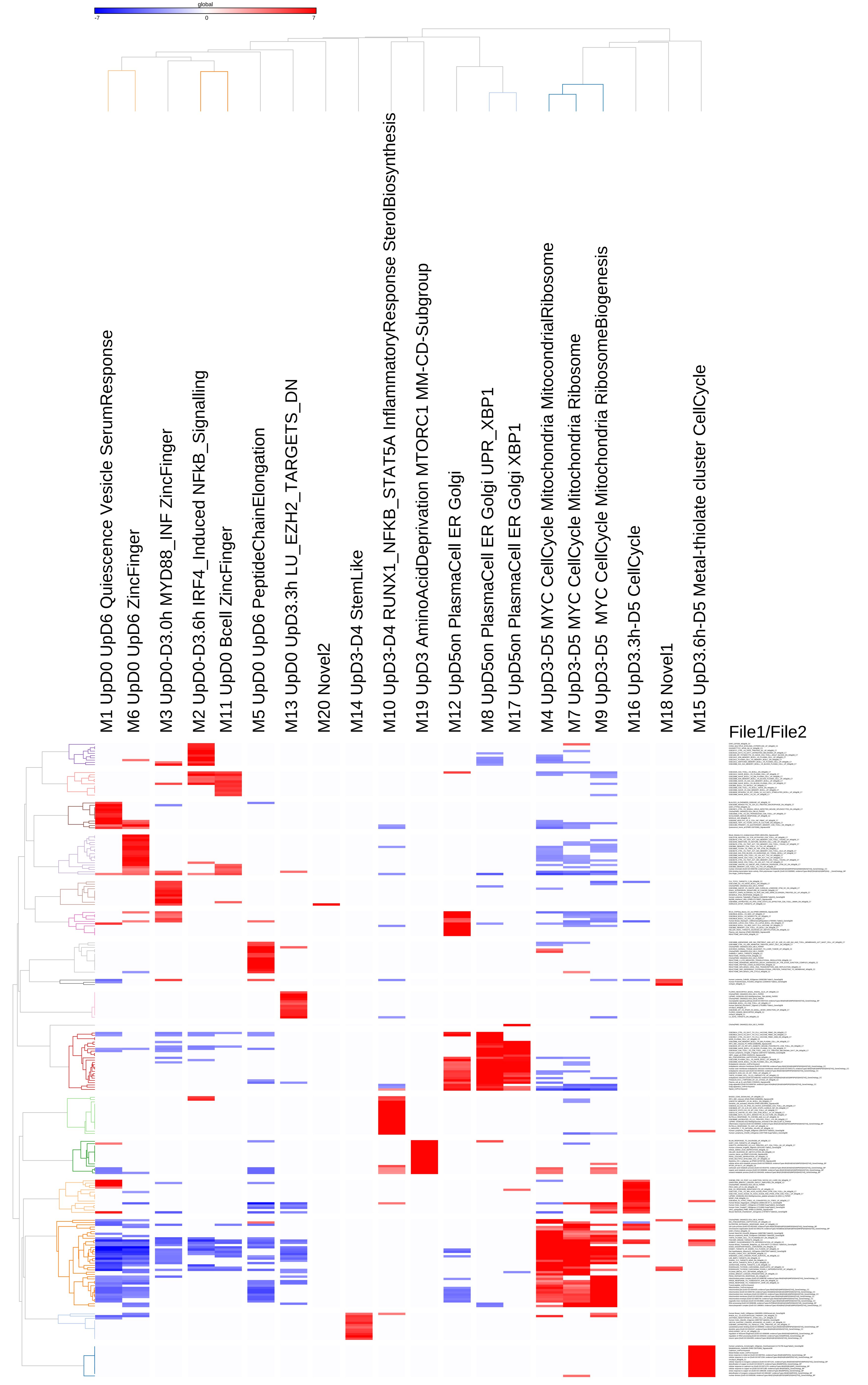

### Supplemental Figure 3

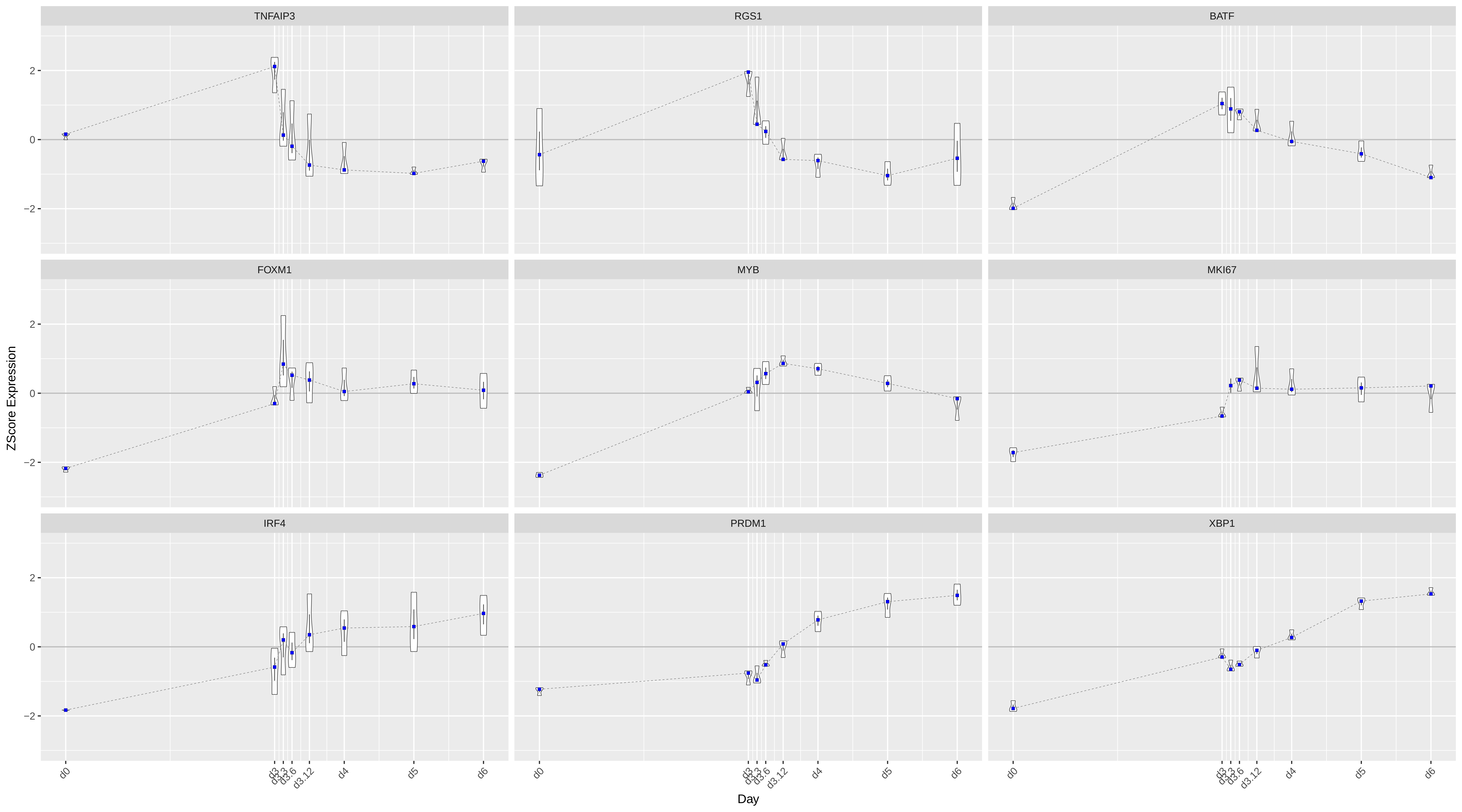

### Supplemental Figure 4

Supplemental Figure 4

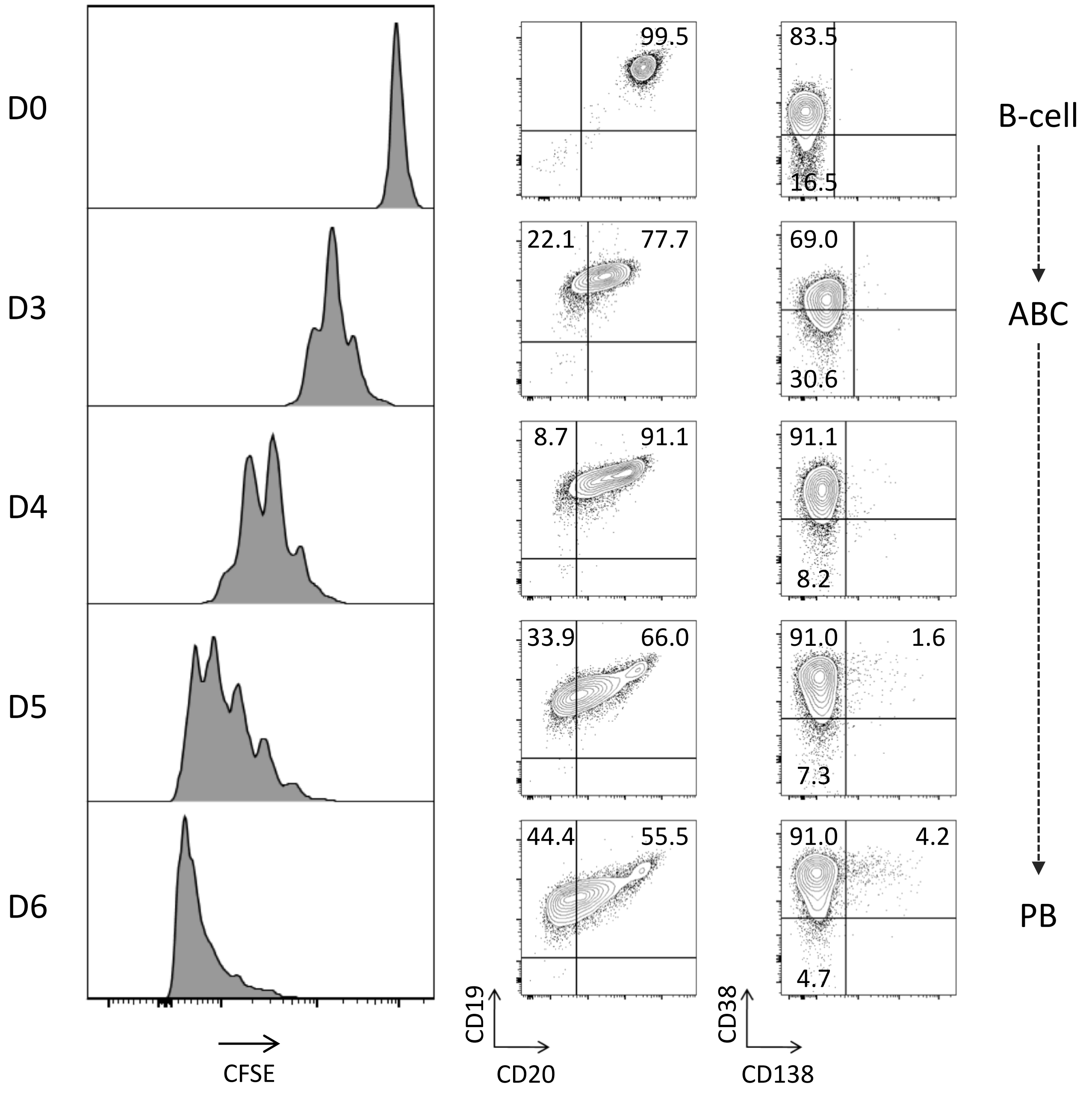

### Supplemental Figure 5

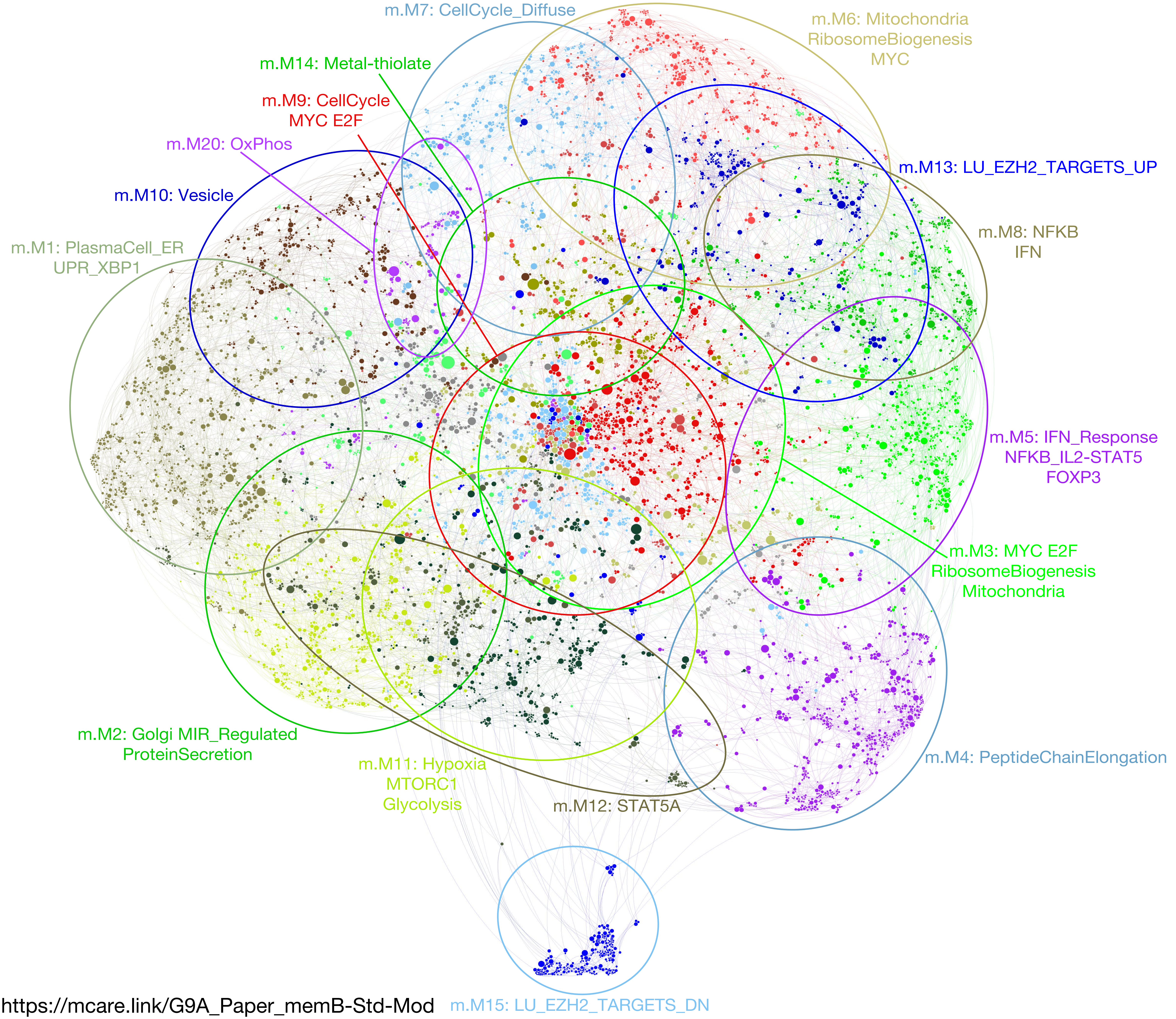

https://mcare.link/G9A_Paper_memB-Std-Mod

### Supplemental Figure 6

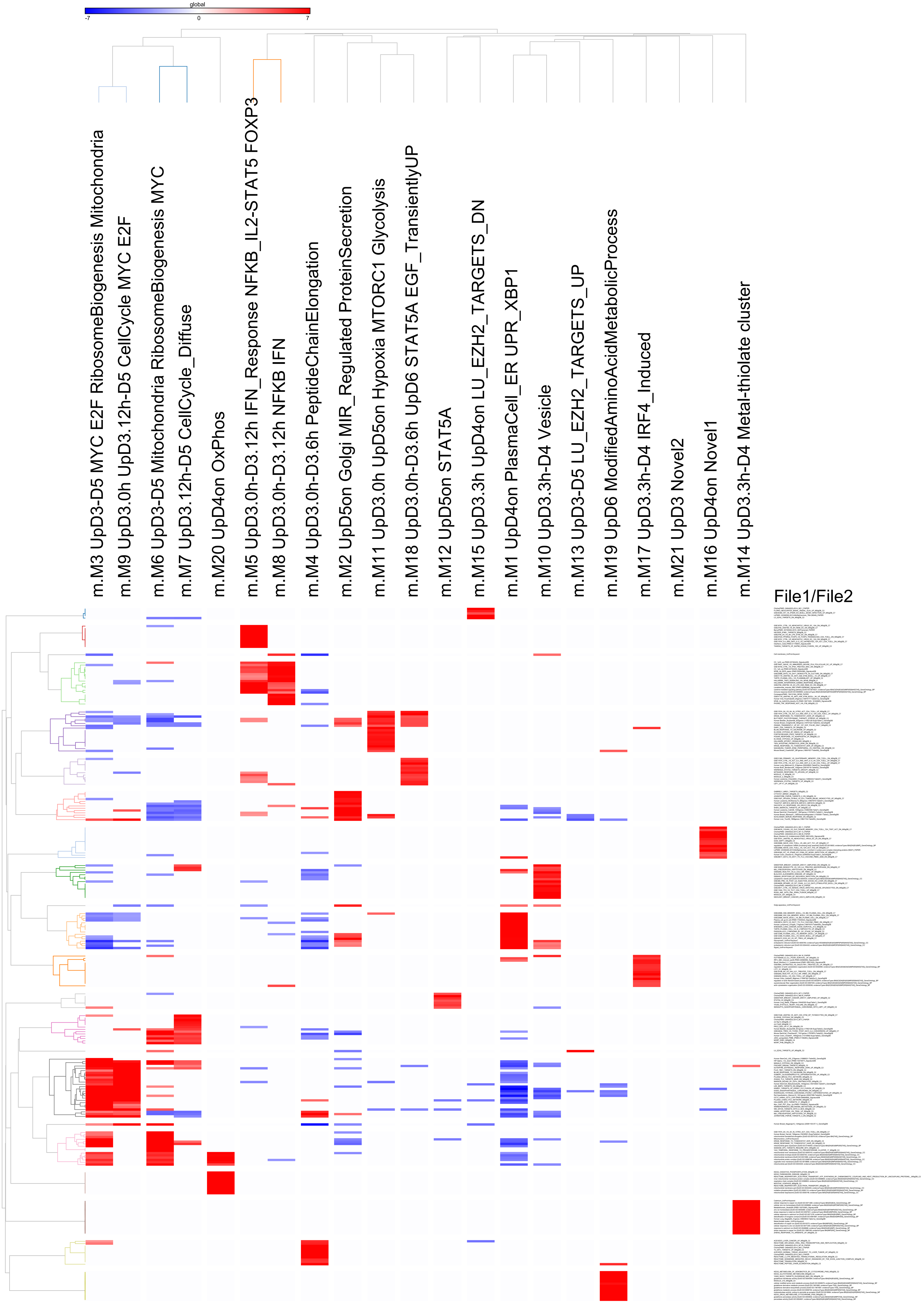

### Supplemental Figure 8

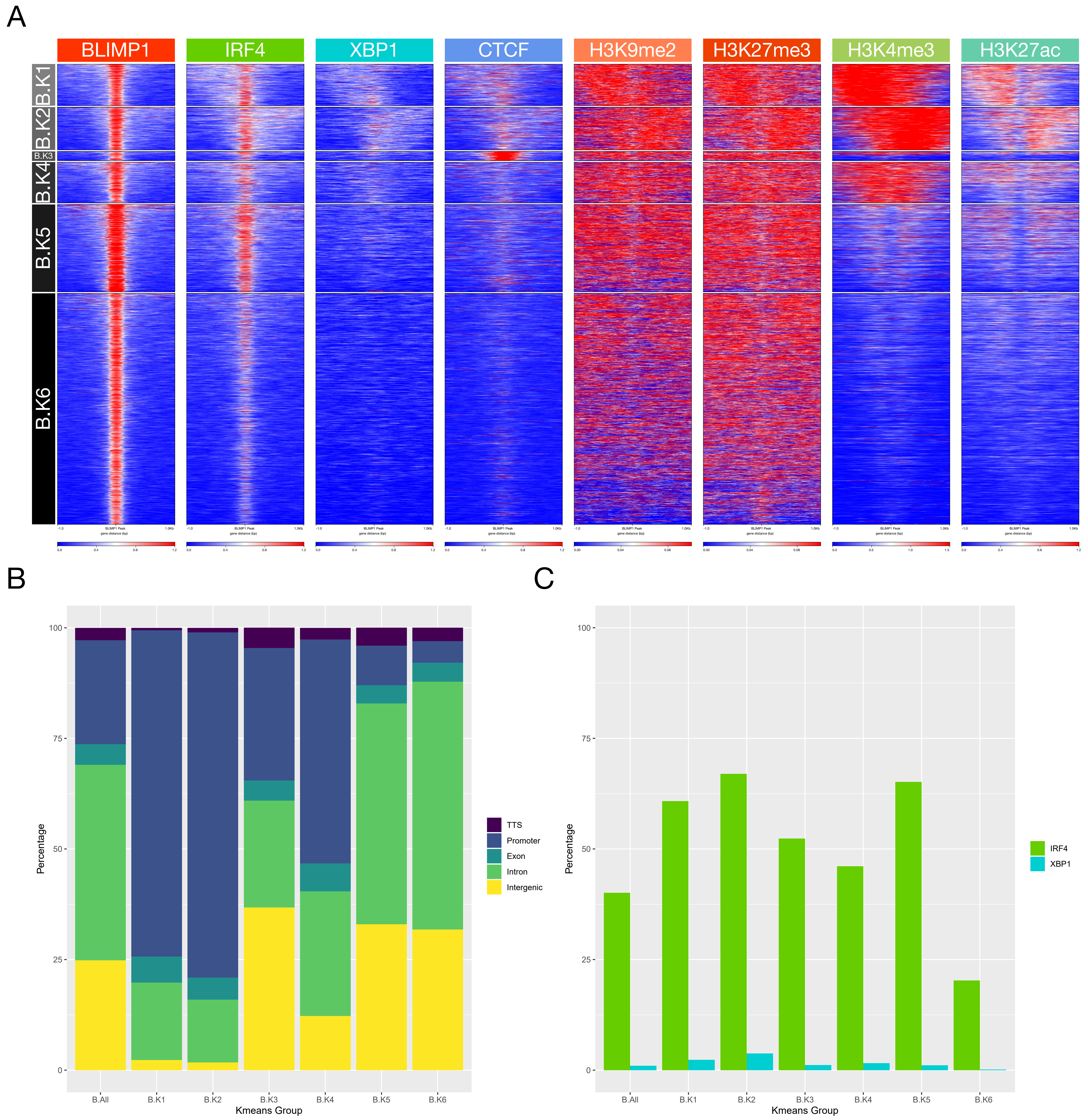

### Supplemental Figure 9

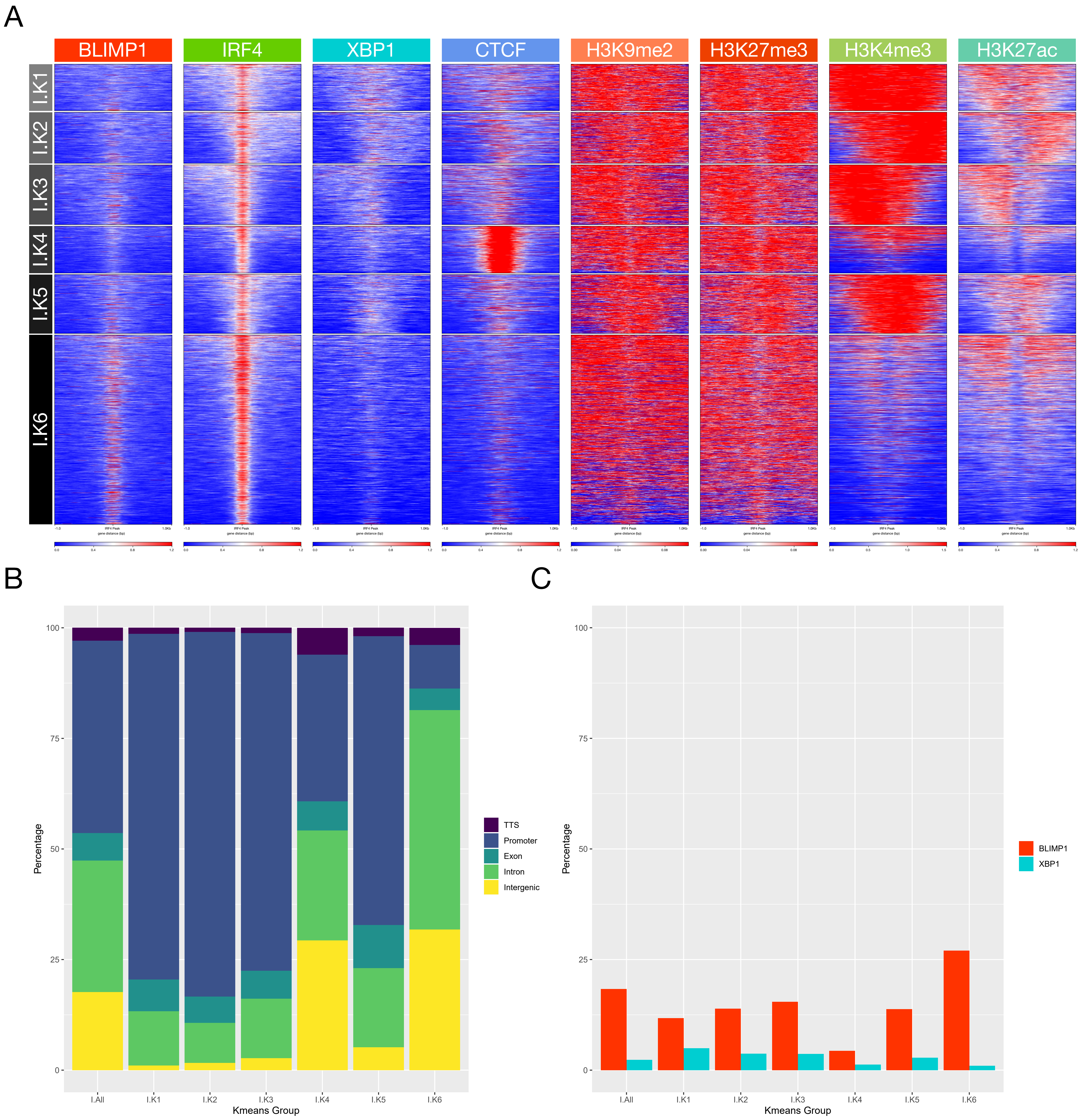

### Supplemental Figure 10

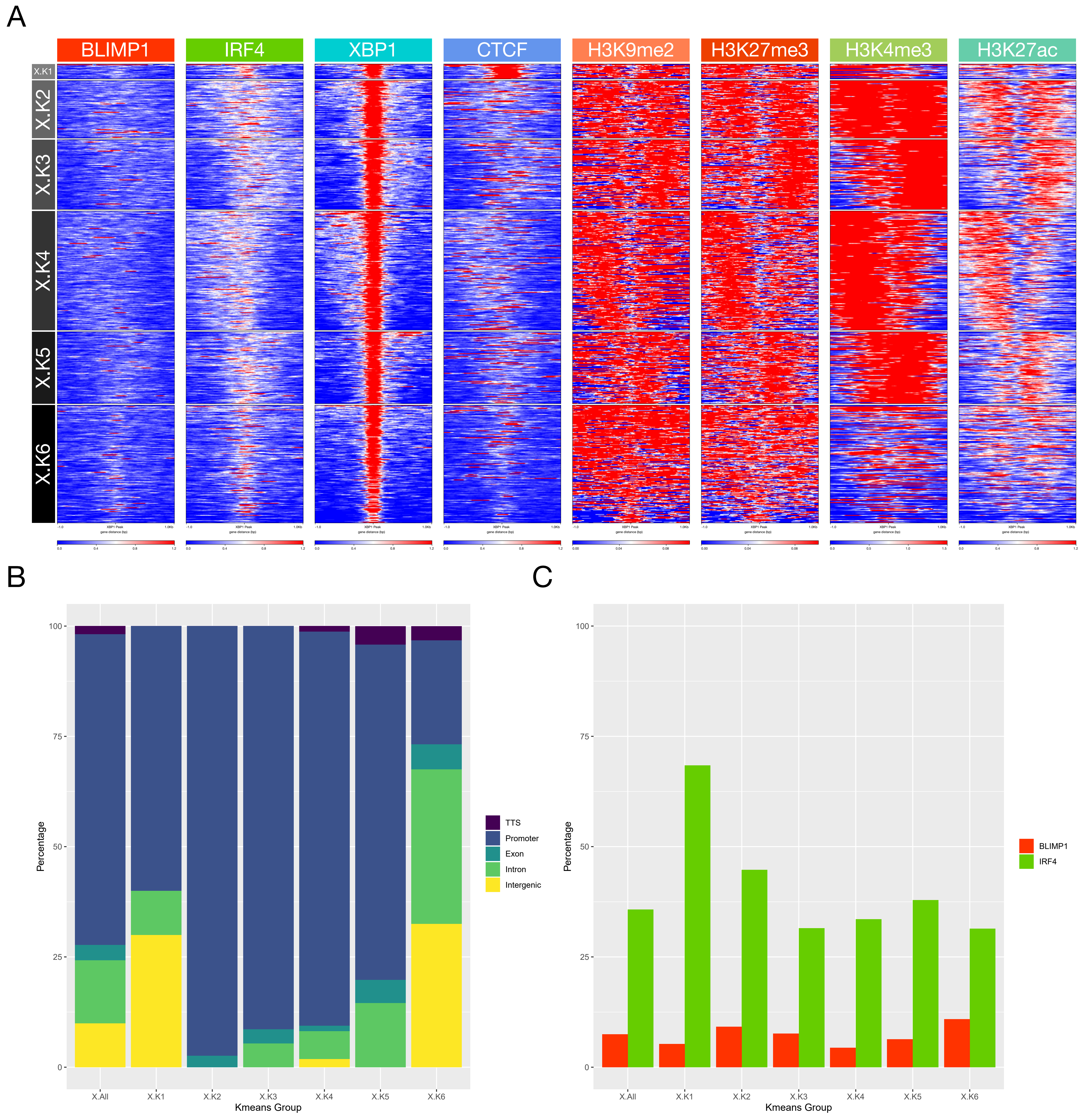

### Supplemental Figure 11

#### Supplemental Figure 11

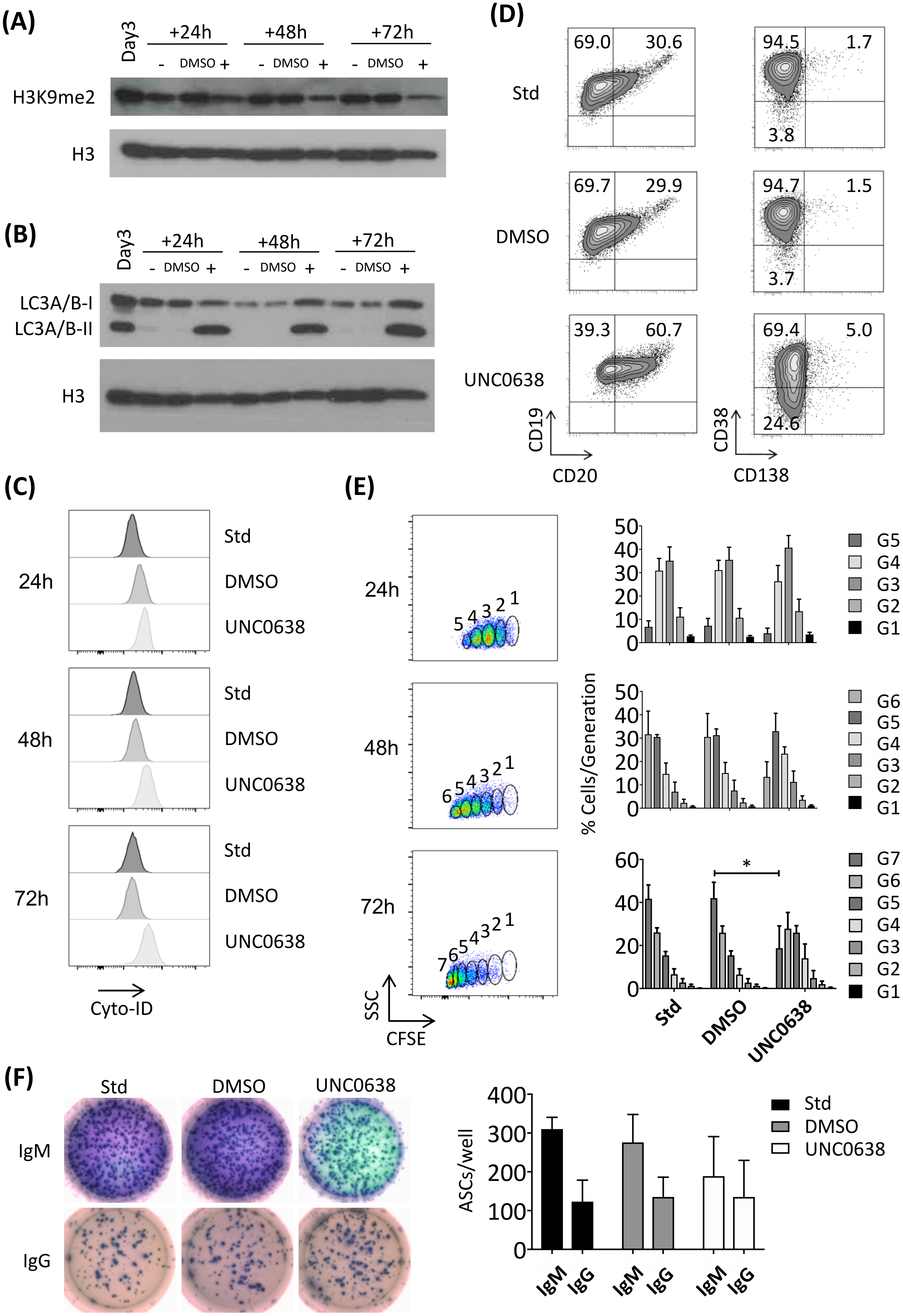
