## Supplemental Figure 7 for "A dichotomy in association of core transcription factors and gene regulation during the activated B-cell to plasmablast transition"

A

### BLIMP1 Only

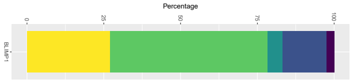


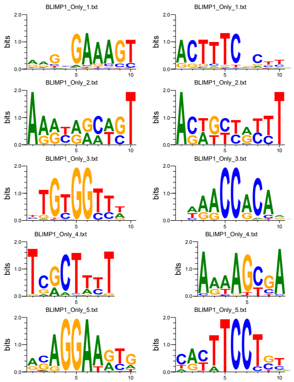

| Forward | Reverse | TF | P-value | %T | %B |
|---|---|---|---|---|---|
| | | PRDM1 | 1e-1179 | 57.03 | 4.13 |
| | | MYF | 1e-173 | 36.21 | 13.24 |
| | | RUNX | 1e-127 | 25.07 | 8.41 |
| | | IRF1 | 1e-83 | 18.36 | 6.48 |
| | | ETS | 1e-62 | 18.70 | 7.90 |

### IRF4 Only

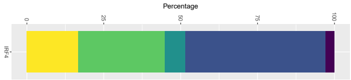


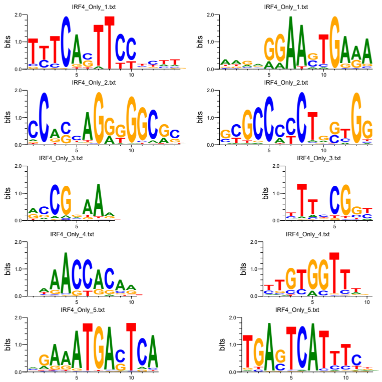

| Forward | Reverse | TF | P-value | %T | %B |
|---|---|---|---|---|---|
| | | ETS:IRF | 1e-701 | 13.72 | 1.09 |
| | | BORIS/CTCF | 1e-319 | 9.85 | 1.53 |
| | | ELK4 | 1e-169 | 63.45 | 46.92 |
| | | RUNX | 1e-155 | 23.20 | 11.82 |
| | | AICE | 1e-96 | 3.31 | 0.57 |

### BLIMP1 & IRF4

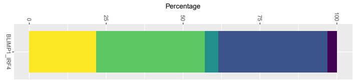


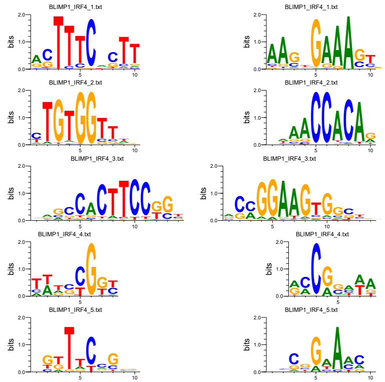

| Forward | Reverse | TF | P-value | %T | %B |
|---|---|---|---|---|---|
| | | PRDM1 | 1e-422 | 51.25 | 10.06 |
| | | RUNX | 1e-95 | 31.71 | 12.82 |
| | | ETS | 1e-87 | 20.92 | 6.60 |
| | | ELK4 | 1e-61 | 47.15 | 28.57 |
| | | ELK4 | 1e-55 | 20.53 | 8.55 |

B

### CTCF

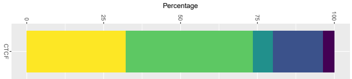


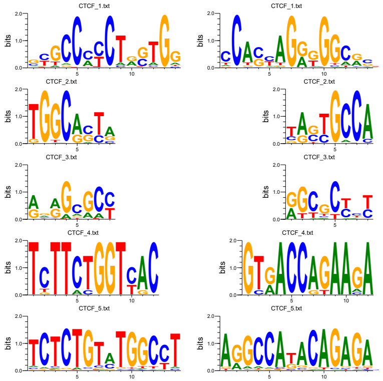

| Forward | Reverse | TF | P-value | %T | %B |
|---|---|---|---|---|---|
| | | BORIS/CTCF | 1e-22756 | 47.21 | 2.08 |
| | | NFIX | 1e-1859 | 56.46 | 35.21 |
| | | BREu | 1e-1121 | 67.17 | 50.56 |
| | | ETS | 1e-393 | 0.45 | 0.00 |
| | | RAP1 | 1e-345 | 0.47 | 0.01 |

C

### XBP1

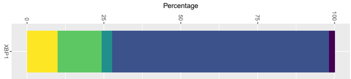


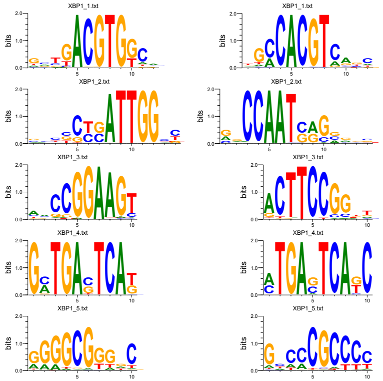

| Forward | Reverse | TF | P-value | %T | %B |
|---|---|---|---|---|---|
| | | XBP1 | 1e-250 | 35.71 | 1.15 |
| | | NFY | 1e-41 | 20.60 | 4.87 |
| | | ETS | 1e-34 | 20.10 | 5.54 |
| | | AP1 | 1e-30 | 8.31 | 0.92 |
| | | SP1 | 1e-29 | 29.57 | 12.02 |

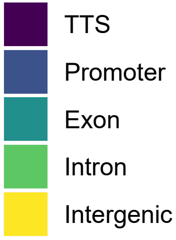
